# Fast and Easy Whole-Brain Network Model Parameter Estimation with Automatic Differentiation

**DOI:** 10.1101/2025.11.18.689003

**Authors:** Marius Pille, Leon Martin, Emilius Richter, Dionysios Perdikis, Michael Schirner, Petra Ritter

## Abstract

Personalized brain modeling at clinically relevant scales requires integrating biophysical models with empirical neuroimaging data, yet high-dimensional parameter estimation in whole-brain network models remains computationally prohibitive. We present TVB-Optim, an open-source Python library providing a general and extensible framework for gradient-based optimization of brain network models build on JAX. Leveraging automatic differentiation, we demonstrate direct optimization of thousands of parameters, from global coupling (N=2) to regional dynamics (N=168) to full structural connectivity matrices (N=14,028), across functional MRI and magnetoencephalography data. Forward simulations achieve 10× CPU speedup over reference implementations and scale efficiently to GPU and multi-device configurations. We establish best practices through documented workflows that combine coarse parameter space exploration with gradient refinement, often yielding superior solutions faster than gradient methods alone. By bridging mechanistic neuroscience with modern machine learning infrastructure, TVB-Optim enables large-scale personalized brain models, bringing computational neuroscience closer to clinical translation.

## 1 Introduction

To understand brain complexity, dynamical systems modeling has proven tremendously effective [1–5], with applications ranging from microscopic neuron circuits [6,7] through mesoscopic systems [8,9] to whole-brain perspectives [10,11]. This mechanistic approach builds understanding from single neurons to larger circuits, integrating prior physiological knowledge as needed. In contrast, data-driven approaches using state-of-the-art Machine Learning (ML) models like deep neural networks or transformers [12,13] require large training datasets. In human medical research, acquiring sufficiently large datasets for classical ML approaches poses severe technical and ethical challenges [14], fundamentally limiting their clinical application. Furthermore, unlike mechanistic modeling that embeds testable hypotheses, black-box models fit data without ensuring biological plausibility. Recent work demonstrates their internal reasoning processes can be fundamentally disconnected from mechanistic logic [15]. Our approach bridges these paradigms by applying the computational backbone of modern ML, automatic differentiation (Automatic Differentiation (AD)) [16], to mechanistic brain models. AD systematically computes derivatives of arbitrary computational graphs with machine precision, enabling efficient gradient-based optimization [17] that addresses the curse of dimensionality. This combination maintains scientific interpretability while enabling effective parameter inference from limited clinical data.

A Brain Network Model (BNM), also termed *Virtual Brain* model [18], *Connectome-Based Neural Mass Model* [19], or *Large-Scale Brain Network Model* [10], simulates large-scale neural dynamics by coupling local neuronal population dynamics across brain regions according to structural connectivity. These models mathematically represent stochastic delay differential equations where neural mass models describing regional population behavior are coupled through delayed connections reflecting white matter pathways. Observation models transform simulated activity into empirically comparable quantities such as Functional Magnetic Resonance Imaging (fMRI) Blood-Oxygenation-Level–Dependent imaging (BOLD) or Electroencephalography (EEG)/Magnetoencephalography (MEG) signals.

Existing BNM simulation tools face tradeoffs between speed, usability, and flexibility that hinder clinical translation [20]. While platforms like The Virtual Brain (TVB) [10,18] provide comprehensive environments, performance-optimized implementations leverage specialized languages [21,22], and methodological innovations address specific inference challenges [23–25], these tools often sacrifice generality, accessibility, or require substantial expertise. Moreover, existing tools lack the integration of modern automatic differentiation with general-purpose BNM frameworks, preventing scalable gradient-based optimization of the thousands of parameters required for personalized modeling. We introduce TVB-Optim, a comprehensive Python library that addresses these gaps by generating fast, differentiable implementations from high-level BNM specifications. Leveraging JAX [26], a numerical library with automatic differentiation capabilities, we present a framework for *Differentiable Brain Dynamics* that embed anatomical, physiological, and biophysical knowledge in model structure and parameters while enabling efficient gradient-based optimization. Integrating with The Virtual Brain Ontology (TVB-O) [27] a structured knowledge base for brain network modelling, the library scales from CPU to GPU clusters, enables optimization of global to connectivity-scale parameters, and provides documented workflows for fMRI, EEG and MEG modeling. This approach validates methods successfully applied to biophysical neuron models [28], bringing differentiable programming to whole-brain network modeling while preserving physiological interpretability.

## 2 Results

TVB-Optim implements a four-step workflow for parameter estimation in brain network models (Figure 1). First, models are defined, either directly or within TVB-O, a convenient knowledge base. This step combines structural connectivity with neural mass model specifications into a networked dynamical system. Second, simulations generate observables, which are compared to empirical data via differentiable loss functions. Third, parallel parameter sweeps map the dynamical landscape, revealing parameter sensitivities and optimal regimes. Fourth, gradient-based optimization refines parameters using automatic differentiation. This workflow supports iterative refinement by progressively expanding the parameter space from global parameters (scalar values identical across all brain regions) to regional parameters (brain-region specific values) to connectivity parameters (modifying individual connections among regions) as illustrated in Figure 7. A key advantage of reverse-mode automatic differentiation is that gradient computation costs remain nearly constant regardless of parameter count *N*_*θ*_, enabling optimization of thousands of parameters.

**Figure 1.**
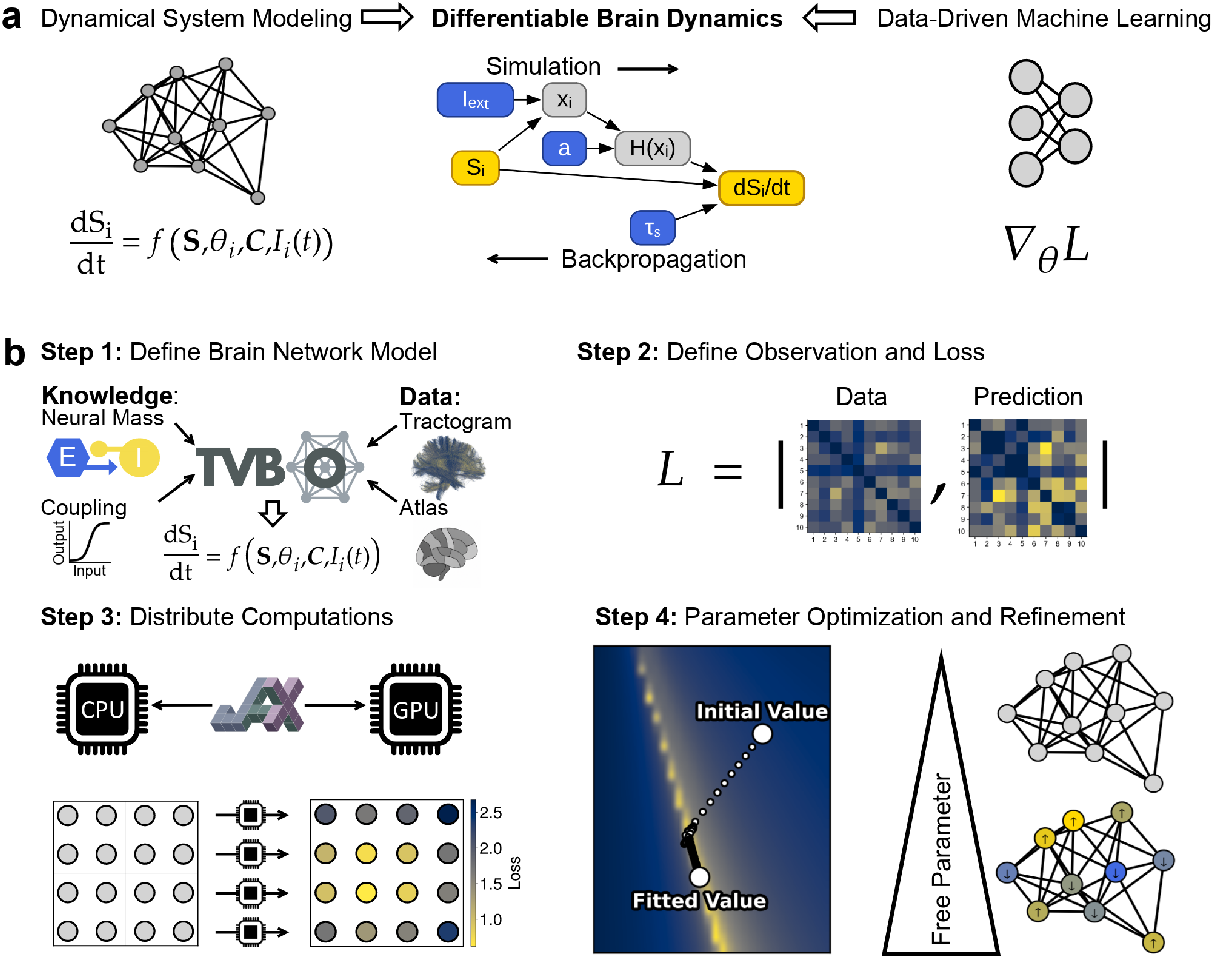
Differentiable brain dynamics framework for parameter estimation in brain network models. **Panel a** illustrates the conceptual bridge between mechanistic brain modeling and data-driven machine learning approaches through differentiable programming. Traditional mechanistic models embed physiological knowledge but struggle with parameter estimation at scale, while machine learning excels at optimization but lacks interpretability. Our approach combines both by making mechanistic BNMs differentiable through automatic differentiation, enabling scalable parameter inference from limited clinical data while preserving physiological interpretability. **Panel b** presents the structured four-step workflow for parameter optimization implemented in TVB-Optim. **Step 1** begins with BNM definition within TVB-O, a structured knowledge base that provides high-level model specifications. The framework combines subject-specific tractography data with standardized brain atlases to establish network structure via structural connectivity matrices. Local dynamics at each node are defined by Neural Mass Model (NMM)s such as the reduced Wong-Wang, Jansen-Rit, or Hopf models, with coupling based on structural connectivity and transmission delays giving rise to the networked dynamical system. Additional forward models transform raw neural activity into observable modalities like fMRI BOLD signals or EEG/MEG recordings, bridging simulation and empirical measurement. **Step 2** illustrates post-processing of simulated time series into specific observables such as functional connectivity matrices, power spectra, or functional connectivity dynamics that can be compared to empirical target data via differentiable loss functions. **Step 3** leverages JAX’s device-agnostic architecture to seamlessly distribute computations across CPUs, GPUs, and TPU clusters, enabling rapid parallel parameter sweeps that map the model’s dynamical landscape. This exploration phase reveals parameter sensitivities, identifies dynamical regimes, and provides informed initialization for subsequent optimization. **Step 4** demonstrates how this landscape knowledge guides automatic differentiation-based optimization of BNM parameters. The workflow supports iterative refinement by progressively expanding the free parameter space from global parameters identical across all regions (N=2-10) to regional parameters allowing heterogeneity (N=84-168) to connectivity-scale parameters modifying individual connections (N>10,000). Additional loss terms can be incorporated at each stage to constrain solutions with multimodal data. A complete code example implementing this workflow is provided in Supplements Figure 13.

We demonstrate these capabilities through progressively complex tasks. Starting with global parameter optimization (*N*_*θ*_ = 2) for resting-state fMRI using the reduced Wong-Wang model, we advance to regional heterogeneity (*N*_*θ*_ = 168) in MEG modeling with the Jansen-Rit model, and ultimately to connectivity-scale optimization (*N*_*θ*_ = 14, 028) with a two-population reduced Wong-Wang model. Each employs different neural mass models suited to the imaging modality. Complete executable code for the first experiment appears in Supplements Figure 13, with all implementations in the online documentation (https://github.com/virtual-twin/tvboptim).

### 2.1 Global Parameters (*N*_*θ*_ = 2) - Exploring and Optimizing BOLD Dynamics

We demonstrate the workflow with global parameter optimization, fitting *N*_*θ*_ = 2 parameters of the reduced Wong-Wang model [29–31] to resting-state BOLD Functional Connectivity (FC) and Functional Connectivity Dynamics (FCD). This represents the most constrained BNM configuration with identical parameters across all brain regions.

Systematic parameter space exploration on a 32 × 32 grid revealed a complex loss landscape with ridge-shaped optima and multiple local minima (Figure 2 panels **a-b**). Using the ADAM optimizer, we compared two optimization strategies, random initialization versus exploration-guided initialization. Random starts (*N* = 100 runs) frequently converged to local optima despite full convergence (Figure 2 panel **c**, yellow). Initialization from the top 1% of explored parameters consistently achieved superior solutions (panel **c**, blue).

**Figure 2.**
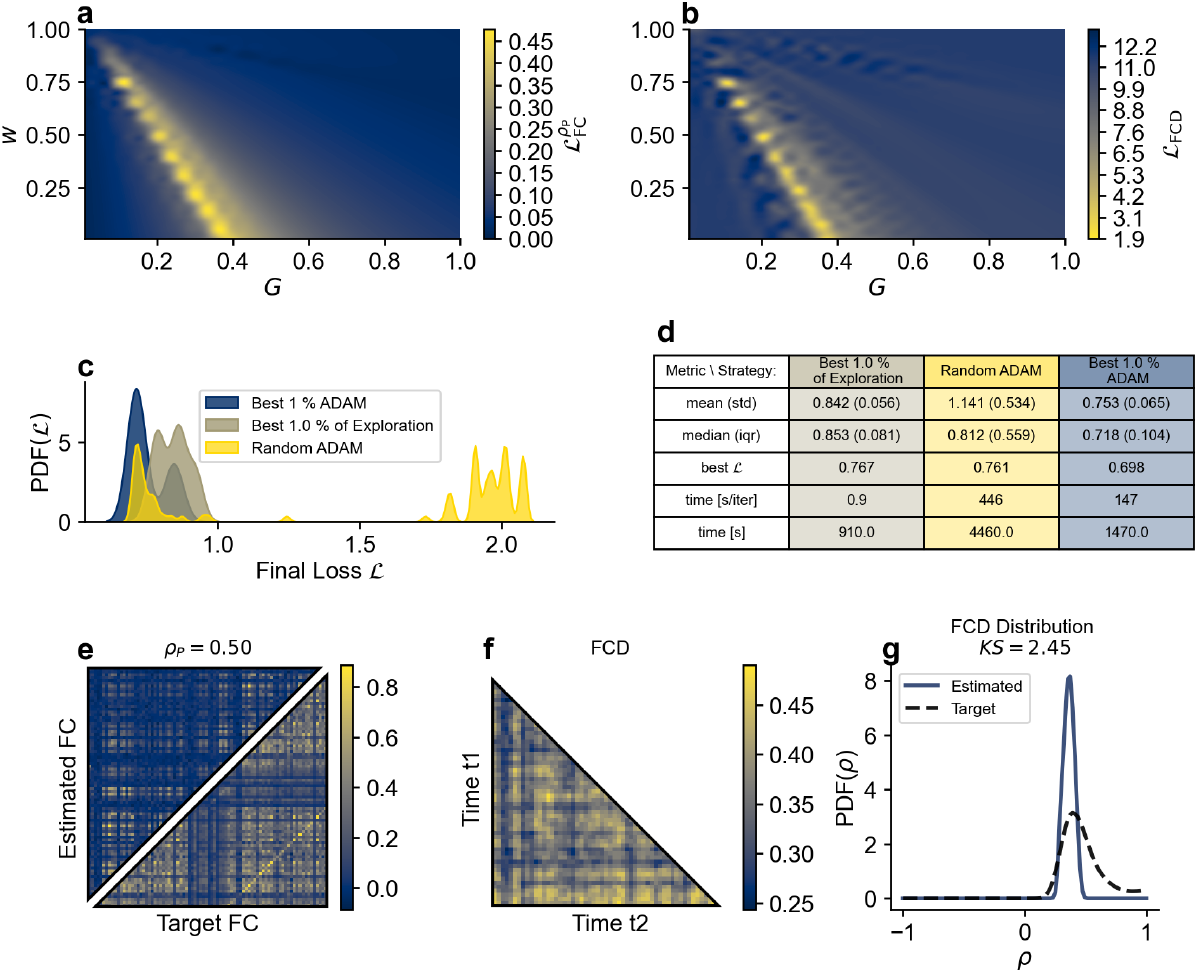
Global parameter optimization workflow combining exploration and gradient-based refinement. We optimized global coupling strength *G* and excitatory recurrence *w* of the reduced Wong-Wang model to match resting-state BOLD FC and FCD from Human Connectome Project data. **Top row** presents grid exploration results (*N* = 1024 parameter combinations on a 32 × 32 grid spanning [0, 1] for both parameters). Panel **a** shows FC correlation loss 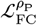 revealing an extended ridge-shaped optimum with negative correlation between *w* and *G*. Panel **b** displays Kolmogorov-Smirnov (KS) statistic for FCD distribution loss ℒ_FCD_, introducing local optima parallel to the ridge that complicate direct optimization. The combined loss function was 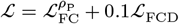. Each exploration simulation required 0.9 seconds, totaling 910 seconds for the complete grid. **Middle row** compares optimization strategies. Panel **c** shows distributions of final losses after 100 optimization iterations with different initialization strategies: ADAM optimizer starting from random initial values (yellow, learning rate *α* = 0.05, *N* = 100 runs), best 1% from exploration without further optimization (tan), and ADAM starting from best 1% of explored parameters (blue, learning rate *α* = 0.001, 10 settings with 10 repetitions each). Random initialization frequently converged to local optima (multimodal distribution) despite full convergence. Initialization from exploration results achieved consistently better solutions. Panel **d** presents timing analysis showing exploration-guided optimization (2380 seconds total: 910s exploration + 1470s optimization) was both more effective and more efficient than random initialization alone (4460 seconds). **Bottom row** shows the best solution achieving FC correlation *ρ*_P_ = 0.50 and FCD KS statistic *D*_KS_ = 2.45. Panel **e** displays estimated versus target FC matrices, panel **f** shows temporal FCD evolution, and panel **g** compares estimated and target FCD distributions. This demonstrates that systematic parameter space exploration is essential for effective optimization in complex loss landscapes.

Crucially, combined exploration-optimization proved both more effective and more efficient. It achieved better solutions while requiring only 2380 seconds total (910s exploration + 1470s optimization) compared to 4460 seconds for random initialization that yielded inferior results (panel **d**). This demonstrates that initial parameter space exploration is essential for efficient optimization in complex loss landscapes. The best solution achieved *ρ*_P_ = 0.50 for FC and *D*_KS_ = 2.45 for FCD (panels **e-g**).

### 2.2 Regional Parameters (*N*_*θ*_ = 168) - Optimizing MEG Peak Frequency Gradient

Scaling from *N*_*θ*_ = 2 to *N*_*θ*_ = 168 parameters (2 parameters × 84 regions), we model the spatial gradient of peak frequencies observed in MEG data (cf. Section 4.2), a fundamental organizational principle of cortical dynamics [32] that cannot be captured with global parameters alone. We employ the Jansen-Rit model [33–36], extensively validated for generating realistic alpha-band oscillations.

We optimized regional parameters *a* and *b* (synaptic time constant and inhibitory post-synaptic potential rate) using ADAM with configuration details in Figure 3 caption. Optimization across *N* = 100 independent runs successfully reproduced the target MEG frequency gradient with *r*_*S*_ = 0.98 (Figure 3 panel **b**). Despite parameter degeneracy where *a* and *b* exhibit positive correlation (panels **c-d**), the optimization consistently converged to functionally similar solutions with median pairwise Spearman correlations of *r*_*S*_ = 0.95 (parameter *a*) and *r*_*S*_ = 0.84 (parameter *b*) across runs. This demonstrates that regional heterogeneity enables capturing spatial gradients in neural dynamics while maintaining solution stability.

**Figure 3.**
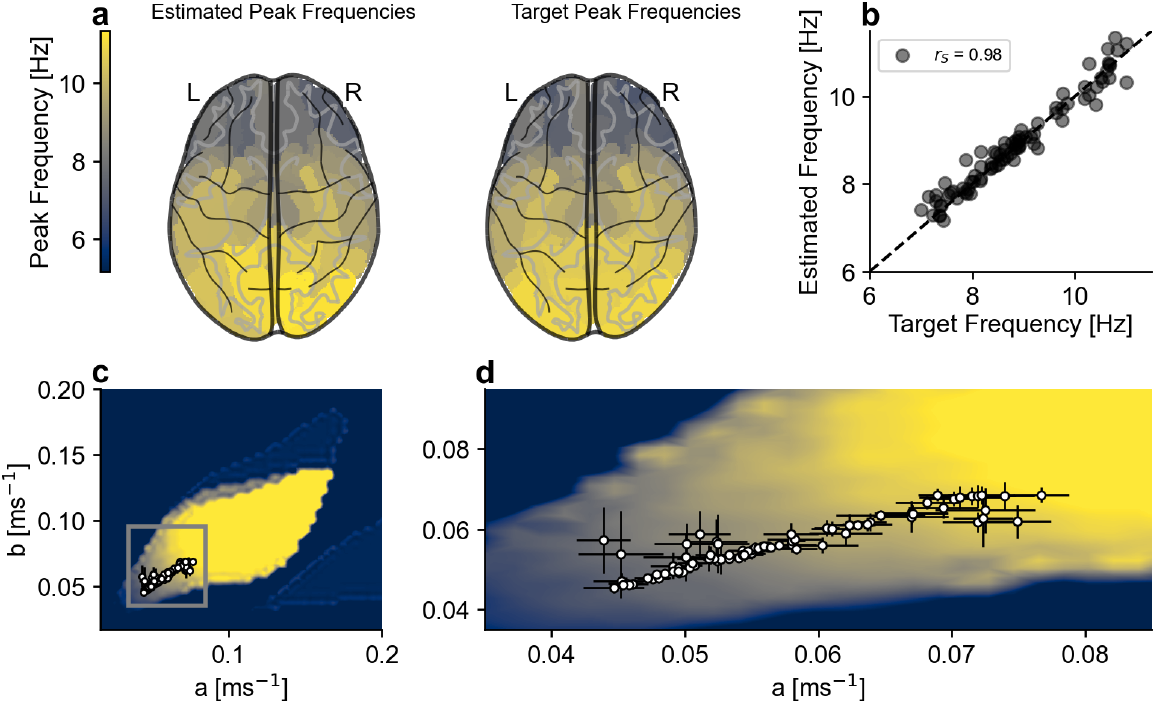
Regional parameter optimization results for MEG peak frequency gradient. We optimized regional parameters *a* and *b* (average synaptic time constant and inhibitory post-synaptic potential rate) of the Jansen-Rit model [33,36] to match the spatial gradient of peak frequencies observed in MEG data (cf. Section 4.2). All 84 regions were initialized with uniform values of *a* = *b* = 0.05 based on prior global exploration (shown as background in panels **c-d**). Global coupling *G* = 15 and external input *I*_*ext*_ = 0.15 were fixed to maintain oscillatory dynamics. Optimization employed the ADAM algorithm with learning rate *α* = 0.001 over 200 iterations, minimizing the loss function specified in Equation 12. Simulations ran for 1000 ms at temporal resolution dt = 1.0 ms, incorporating white noise (standard deviation *σ* = 0.01) via state variable *S*_4_. **Panel a** displays spatial distributions of estimated peak frequencies (left) and target frequencies (right) projected on a glass brain, demonstrating successful reproduction of the anterior-posterior gradient. **Panel b** quantifies the frequency match, plotting estimated versus target peak frequencies for all 84 regions. The strong linear relationship achieves Spearman correlation *r*_*S*_ = 0.98, indicating accurate capture of the spatial gradient. **Panels c-d** reveal parameter degeneracy in the solution space. Parameters *a* and *b* exhibit strong negative correlation across regions, shown against the background of mean peak frequency exploration. Panel **c** presents the full parameter space, while panel **d** zooms into the relevant region. Despite this degeneracy where multiple parameter combinations produce similar dynamics, optimization across *N* = 100 independent runs converged to consistent functional solutions. Median pairwise Spearman correlations across runs were *r*_*S*_ = 0.95 for parameter *a* (IQR: 0.015) and *r*_*S*_ = 0.84 for parameter *b* (IQR: 0.056), demonstrating that the loss landscape effectively constrains the model to reproduce target dynamics while allowing parameter flexibility within a manifold of equivalent solutions.

### 2.3 Connectivity Parameters (*N*_*θ*_ = 14, 028) - Reproducing EI-Tuning

Scaling to *N*_*θ*_ = 14, 028 parameters, we optimize individual connectivity matrix elements, a scale otherwise only tractable with specialized algorithms. We employ a 2-population (excitatory/inhibitory) reduced Wong-Wang model [37,38] incorporating long-range excitation *w*_*LRE*_ and feedforward inhibition *w*_*FFI*_ parameters (cf. Equation 18, Section 4.2.1).

We reimplemented the EI-Tuning algorithm [38] as baseline (see Figure 4 caption for algorithm details). To enable Loss-Based Optimization, we derived a differentiable loss function ℒ_*EI*_ Equation 1 capturing EI-Tuning’s essential components:

**Figure 4.**
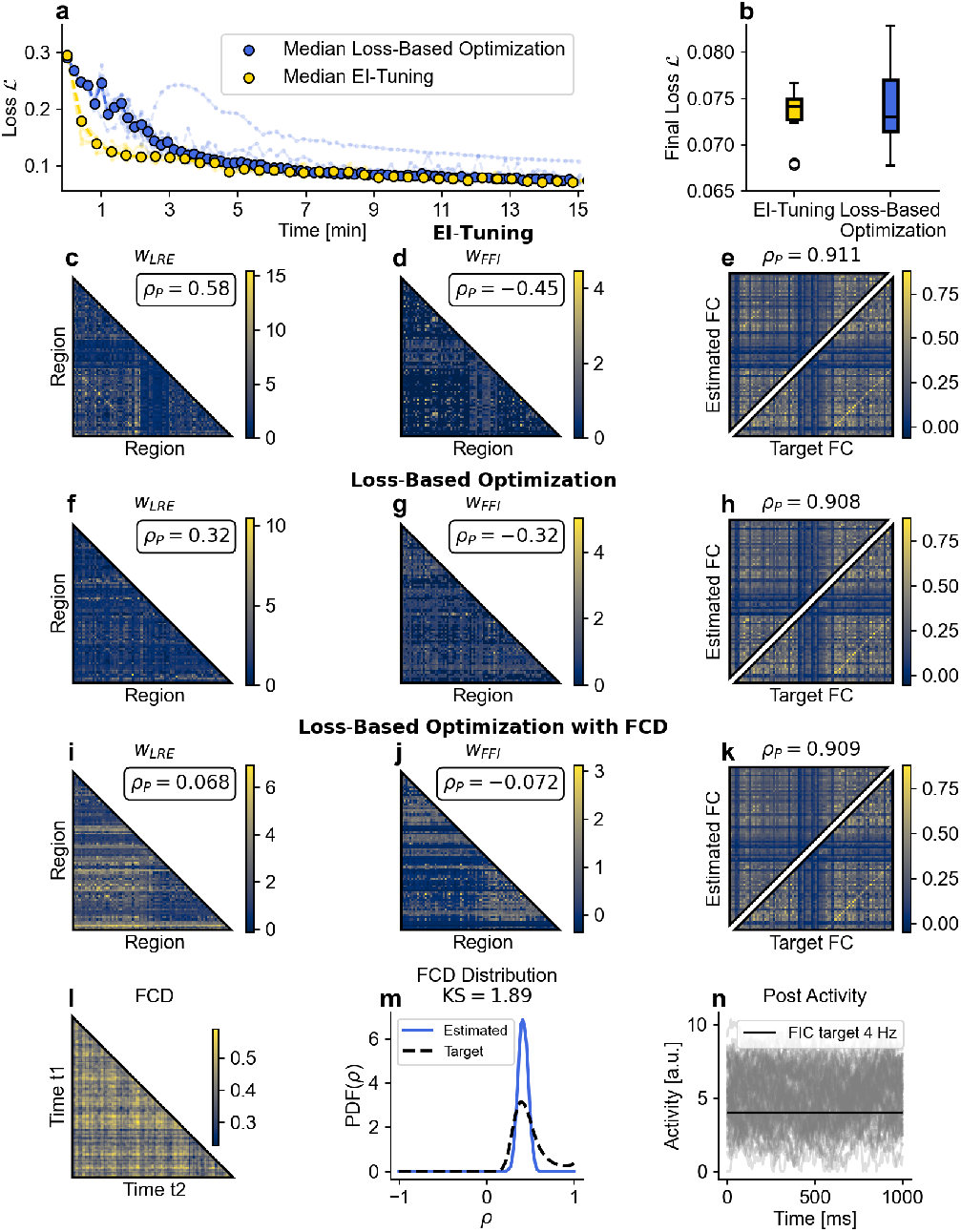
Connectivity-scale parameter optimization comparing EI-Tuning and Loss-Based Optimization. We optimized *N*_*θ*_ = 14028 parameters comprising long-range excitation *w*_*LRE*_ and feedforward inhibition *w*_*FFI*_ connectivity matrices (each containing 6972 parameters from 84×84 connections with self-connections set to 0), plus regional parameters *J*_*i*_ (84 parameters). The 2-population reduced Wong-Wang model [37,38] implements BOLD FC modeling with excitatory/inhibitory balance. **EI-Tuning algorithm** [38]: Iteratively simulates predicted *F C* and updates parameters based on target FC differences. Simultaneously tunes *J*_*i*_ via Feedback Inhibition Control (FIC) [37] to maintain approximately 4 Hz excitatory firing rates. Adaptive strategy increases simulation lengths and decreases learning rates until convergence. **Loss-Based Optimization**: Uses differentiable loss function ℒ_*EI*_ combining FC RMSE and FIC constraint (cf. Equation 1, Equation 2), minimized with ADABelief optimizer (learning rate *α* = 0.2, simulation length *T*_*sim*_ = 10 min). Both approaches ran for 15 minutes wall time across *N* = 10 repetitions. **Panel a** displays loss trajectories over time for all 10 repetitions (thin lines) and their medians (bold) for EI-Tuning (yellow) and Loss-Based Optimization (blue). EI-Tuning demonstrates faster initial convergence and smoother trajectories. **Panel b** compares final loss distributions via boxplots, showing EI-Tuning’s more consistent performance with lower variability. **Panels c-e** present EI-Tuning’s best result: optimized *w*_*LRE*_ matrix (**c**) and *w*_*FFI*_ matrix (**d**) with Pearson correlations *ρ*_*P*_ to target FC, and resulting fitted FC (**e**) achieving *ρ*_*p*_ ≈ 0.91. **Panels f-h** show Loss-Based Optimization’s best result with comparable FC correlation but lower parameter-to-FC correlations and higher activity variance. **Panels i-n** demonstrate Loss-Based Optimization’s flexibility by incorporating FCD loss (cf. Equation 3). The resulting *w*_*LRE*_ (**i**) and *w*_*FFI*_ (**j**) matrices show even lower parameter-to-FC correlations, while the fitted FC remains similar (**k**) and additional FCD features appear. Panel **l** shows the FCD matrix, panel **m** shows the FCD distribution comparing target (dashed) and estimated values with the Kolmogorov-Smirnov statistic, and panel **n** shows the mean neural activity distribution around the 4 Hz target.

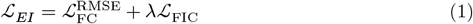

where 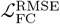 measures the fit to target FC (cf. Equation 10) and ℒ_FIC_ enforces the 4 Hz firing rate constraint,

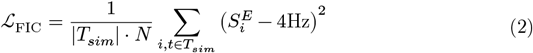

where 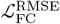 measures FC fit and ℒ_FIC_ enforces 4 Hz firing rates. Parameter constraints like non-negativity are expressed differentiably by transforming *w*_*FFI*_ → |*w*_*FFI*_| and *w*_*LRE*_ → |*w*_*LRE*_| at each simulation start. We performed optimization using ADABelief with *α* = 0.2 and *T*_*sim*_ = 10min (cf. Section 4.4.3).

Both approaches ran for 15 minutes across *N* = 10 repetitions. EI-Tuning showed faster initial convergence (Figure 4 panel **a**) and more consistent performance, while Loss-Based Optimization exhibited larger variability. Both methods achieved comparable results with FC correlation *ρ*_*p*_ ≈ 0.91 and mean activity near the 4 Hz target (panels **c-e** vs. **f-h**). Despite similar metrics, Loss-Based Optimization showed lower correlation between final parameters and target FC, and higher variance in activity patterns. This demonstrates that gradient-based optimization remains viable even at 14,028 parameters, matching specialized algorithms and enabling personalized brain modeling at unprecedented detail.

#### 2.3.1 Adding FCD to the EI-Tuning approach

A key advantage of our differentiable loss-based approach becomes apparent when extending optimization constraints. While specialized algorithms like EI-Tuning rely on neurobiologically-informed mechanisms that are difficult to modify without reimplementation, our formulation enables straightforward integration of arbitrary differentiable constraints. This represents a general capability applicable to any optimization scenario. We demonstrate this flexibility by adding the FCD distribution loss (cf. Equation 11):

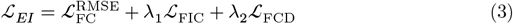

The optimizer successfully finds solutions with comparable FC correlation (Figure 4 panel **k**). However, the additional FCD constraint pushes optimization toward different parameter regimes, with lower parameter-to-FC correlations (panels **i-j**).

### 2.4 Multimodal Data (*N*_*θ*_ = 169) - Combining temporal scales with MEG and fMRI Data

Our differentiable approach enables simultaneous optimization against multiple imaging modalities operating at different timescales, addressing parameter degeneracy by constraining models with complementary data sources [39]. We demonstrate this using the Hopf model [40] (cf. Section 9.1.3), optimizing *N*_*θ*_ = 169 parameters, natural frequency *ω* and bifurcation parameter *a* for each of the 84 regions, plus global coupling *G*. The combined loss simultaneously fits slow BOLD FC and fast MEG peak frequency gradients (optimization details in Figure 5 caption).

**Figure 5.**
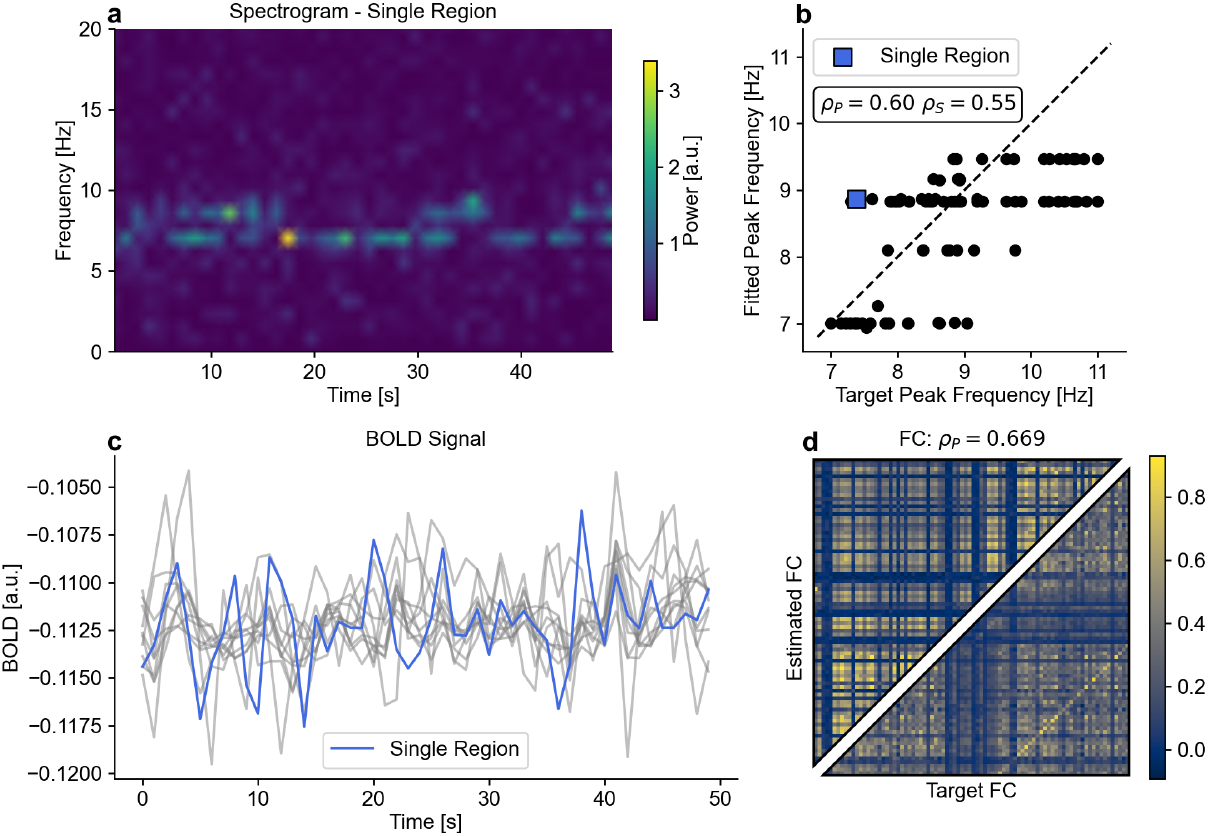
Multimodal and multiscale optimization results combining MEG and fMRI data. We optimized the Hopf model [40] with difference coupling (cf. Section 9.1.3) to simultaneously fit slow BOLD FC and fast MEG peak frequency gradients. The model comprises *N*_*θ*_ = 169 parameters: natural frequency *ω* and bifurcation parameter *a* for each of 84 regions (168 parameters), plus global coupling strength *G*. The combined loss function sums individual modality losses, conditioning simulated BOLD on target FC while matching raw neural activity to regional peak frequencies. Balancing these different timescales required *T*_*sim*_ = 1 min simulations at *dt* = 1 ms resolution. Optimization employed RMSProp with learning rate *α* = 0.0005 over 100 iterations. **Panel a** displays the spectrogram of neural activity from a single representative brain region (highlighted in blue across all panels), showing how the region’s frequency content evolves over time. The spectrogram reveals that regions dynamically switch between different frequency bands and alternately synchronize and desynchronize with other regions, a key property for realistic brain dynamics. **Panel b** presents the fit of the frequency gradient across all 84 brain regions. Synchronization is visible as frequencies cluster into discrete horizontal levels, with multiple regions sharing similar frequencies. The highlighted single region (blue square) corresponds to the same region shown in panel **a.** Pearson and Spearman correlations quantify the accuracy of frequency gradient fitting, achieving *ρ*_*P*_ = 0.60 for the peak frequency match. Compared to FC-only optimization (not shown), the additional frequency constraint reduces inter-regional synchronization, with regions forming smaller synchronized clusters rather than global synchrony. **Panel c** displays the BOLD signal time courses, with the highlighted single region (blue) overlaid on 10 other representative regions (grey), demonstrating realistic temporal dynamics. **Panel d** compares target and estimated FC matrices, achieving correlation *ρ*_*P*_ = 0.669, comparable to FC-only optimization while simultaneously satisfying the frequency gradient constraint. This demonstrates that multimodal constraints can mitigate parameter degeneracy by providing complementary information across timescales.

Multimodal constraints shaped the solution space differently than single-modality optimization (Figure 5). The additional frequency constraint reduced global synchronization, with regions forming smaller synchronized clusters (panel **b**). The model successfully captured both targets with peak frequency correlation *ρ*_*P*_ = 0.60 and FC correlation *ρ*_*P*_ = 0.669, comparable to FC-only optimization.

### 2.5 Performance

We benchmarked TVB-Optim across computational dimensions critical for large-scale parameter optimization workflows (Figure 6). Forward simulation speed was evaluated against the established TVB implementation. Panels **a** and **b** show that our JAX backend achieved 10-50× speedups across all 21 default TVB neural mass models, with performance depending on model complexity and coupling mechanisms. Simpler models reached up to 90× acceleration, while delay-coupled networks maintained 10-25× improvements. Parallel execution efficiency was assessed for parameter space exploration tasks. Panel **d** demonstrates that GPU-based sweeps scaled near-linearly with batch size, while distribution across 4 GPUs provided 3× speedup, indicating efficient multi-device utilization. Most critically for high-dimensional optimization, reverse-mode automatic differentiation maintained nearly constant computational cost regardless of parameter count. Panel **e** shows that while forward-mode differentiation scaled linearly, reverse-mode computation time remained stable whether optimizing 2 or 14,000 parameters. This constant-time gradient property fundamentally enables the connectivity-scale optimizations demonstrated earlier (see Section 4.5.4 and Figure 6 for detailed specifications).

**Figure 6.**
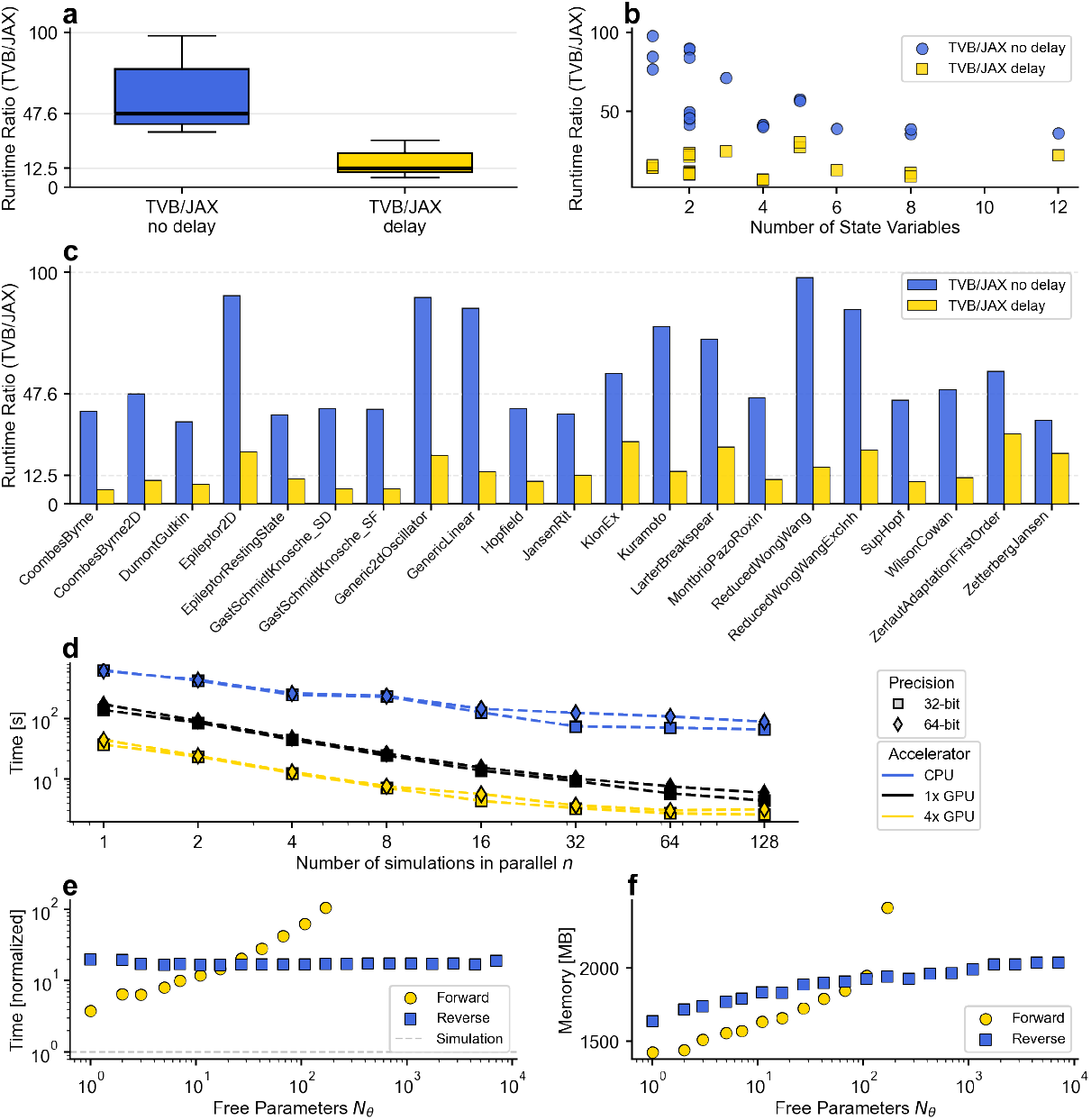
Computational performance evaluation of TVB-Optim. We benchmarked the JAX implementation across factors critical for optimization workflows (see Methods for specifications). Panels **a-b**: Forward simulation performance compared to TVB across all 21 default neural mass models (100,000 timesteps, 84-region networks with stochastic noise). Panel **a** displays runtime ratio distributions (TVB time / JAX time) for simulations with and without axonal transmission delays. The JAX implementation achieved mean speedups of 57.5× without delays (median 47.6×, 10-90th percentiles 38.5-89.3×) and 15.1× with delays (median 12.5×, 10-90th percentiles 6.7-24.5×). Delay handling requires additional memory operations that reduce but do not eliminate the performance advantage. Panel **b** plots these speedup ratios against model complexity (measured by state variable count), demonstrating consistent performance gains across different architectures. Panel **c**: Individual model breakdown comparing delay versus no-delay conditions. Simpler models achieve higher acceleration factors without delays (up to 90×), while delay-enabled simulations show more uniform but lower speedups (10-25×) across all models. Panel **d**: Parallelization efficiency measured as total wall time for *N* = 1024 simulations with varying parallelization *n* (maximum *n* = 128 limited by device memory). CPU performance saturated quickly as thread count increased, while GPU performance scaled nearly linearly with batch size. Distribution across a 4-GPU node provided an additional 3.02× speedup, demonstrating efficient scaling crucial for large parameter space explorations. Panels **e-f**: Automatic differentiation efficiency, the key enabler of high-dimensional optimization. Panel **e** shows computation time normalized to forward simulation cost. Reverse-mode AD time remains nearly constant regardless of parameter count, while forward-mode AD scales linearly with parameter number. This constant-time gradient property fundamentally enables the connectivity-scale optimizations with thousands of parameters demonstrated in Section 2.3. Panel **f** : Memory usage comparison, which favors forward-mode AD below approximately 200 parameters; beyond this threshold, reverse-mode becomes essential despite higher memory overhead. While 32-bit precision improves computational performance, 64-bit precision is required for gradient computations to maintain numerical stability.

## 3 Discussion

TVB-Optim demonstrates that automatic differentiation transforms brain network modeling, enabling optimization of biophysically detailed models at unprecedented scales. This advance addresses a critical bottleneck where mechanistic models remain largely qualitative due to parameter estimation challenges, limiting their clinical utility despite containing rich biological detail. With 10x speedups and gradient-based optimization of thousands of parameters, differentiable brain dynamics bridges mechanistic understanding with data-driven fitting. We discuss implications for computational neuroscience, limitations at dynamical bifurcations, and paths toward clinical translation.

Our performance improvements address a fundamental bottleneck in efficient exploration of complex parameter spaces. Grid search remains dominant for parameter estimation despite exponential scaling (*n*^*D*^ for *D* parameters) that limits exploration to few parameters. Its popularity stems from easy applicability and straightforward interpretability. Despite limitations, exploration provides crucial insights into dynamical landscapes and identifies valid initialization points for optimization. We maximize exploration efficiency while maintaining simplicity. Expressing BNMs in JAX provides robust 10x performance gains over TVB (cf. Figure 6 panel **a**), representing an optimal trade-off between performance and generality. While specialized implementations like fastTVB achieve higher single-simulation speeds, our approach maintains compatibility across all neural mass models while optimizing for gradient computation.

Beyond single-simulation speedups, exploration parallelizes naturally across cores, machines, and accelerators. JAX’s inherent parallelization drastically reduces iteration times (cf. Figure 6 panel **d**) while targeting modern hardware without specialized coding knowledge. Algorithmic improvements complement these gains. Switching from grid search to random sampling or Sobol sequences often yields significantly better results [41]. All schemes evaluate efficiently in parallel. Exploration results integrate seamlessly with simulation-based inference [42], proven powerful for obtaining posterior distributions with limited parameters [43]. Sobol sequences enable direct use for sensitivity analysis [44]. These performance gains make large-scale experiments practical, enabling systematic investigations of model degeneracy, uncertainty quantification, and sensitivity analysis that inform biological interpretation.

While efficient exploration provides a foundation for low-dimensional problems, realistic brain modeling requires refining hundreds to thousands of parameters where traditional approaches fail. Reverse mode AD uniquely addresses this through near-constant computational costs regardless of parameter count (cf. Figure 6 panel **e**), enabling complex loss optimization with Stochastic Gradient Descent (SGD) family algorithms. Our results demonstrate a systematic progression. Gradient-based optimization refines two-parameter grid searches (Section 2.1), transforms global parameters into region-specific ones capturing spatial gradients (Section 2.2), and scales to thousands of connectivity parameters matching specialized algorithms (Section 2.3). Crucially, reformulating the iterative EI-tuning algorithm as a differentiable loss not only validated gradient-based optimization at scale but revealed solution space structure. Different optimization paths yielded comparable performance yet distinct parameter correlations, suggesting underconstrained problems with multiple valid solutions. This observation naturally motivates constraining solution spaces through multiple empirical targets.

Creating physiologically realistic brain models often requires matching multiple empirical observations from different modalities characterized by distinct spatio-temporal scales. TVB-Optim enables straightforward combination of metrics from fMRI, MEG, or EEG through differentiable scalar functions. Where operations like histograms lack inherent differentiability, smooth approximations (e.g., kernel density estimation for FCD) maintain information while enabling optimization. This flexibility proves valuable for constraining underconstrained problems. Adding FCD constraints to EI-tuning eliminated spurious parameter correlations, improving physiological plausibility (Section 2.3). Similarly, simultaneously optimizing for slow fMRI dynamics and fast MEG oscillations acts as physiological regularization, preventing unrealistic synchronization patterns while maintaining empirical fit (Section 2.4). We normalize loss components by maximum values before addition, providing sensible default weights. Beyond technical convenience, integrating multiple empirical targets fundamentally constrains solution spaces toward biologically plausible parameter regimes, demonstrating how differentiable frameworks naturally support multi-modal, multi-scale brain modeling.

While we focused on optimization for point estimates, the gradient infrastructure established here enables a much broader spectrum of inference approaches. The same gradient computations that power optimization directly support Bayesian methods including variational inference and Hamiltonian Monte Carlo, enabling researchers to transition seamlessly from point estimates to full posterior distributions as their scientific or clinical questions demand (see Section 9.6 for detailed discussion).

Despite these capabilities, fundamental limitations arise at dynamical bifurcations where many brain models operate optimally (detailed technical discussion in Supplements Section 9.5). Brain network models frequently achieve optimal fits at or near bifurcation points where system dynamics change qualitatively [45,46]. Near these critical points, small parameter changes induce large dynamical shifts, producing extremely large gradients that can destabilize optimization [47]. Gradient clipping addresses this by bounding gradient magnitudes (Section 2.1). More severe instabilities emerge when optimal fits occur where the maximum Lyapunov exponent becomes positive, indicating chaotic dynamics. Standard automatic differentiation becomes unsuitable when simulation time exceeds the Lyapunov time. Solutions include gradient flossing [48], which constrains Lyapunov exponents as differentiable terms, or shadowing methods [49] that compute meaningful gradients in chaotic regimes. Both approaches face practical limitations. Gradient flossing requires careful parameter selection given physiological meaning of BNM parameters, while shadowing methods scale poorly with network size and require adaptations for delayed systems [50]. Beyond these dynamical challenges, current fixed step size integration prevents gradient-based optimization of delay parameters due to non-smooth derivatives from discrete history indexing. Continuous, differentiable interpolation would enable delay optimization while supporting adaptive time stepping [51].

These advances establish TVB-Optim as a comprehensive framework for parameter estimation in brain network models, combining automatic differentiation with biological detail. With 10x performance improvements and optimization of thousands of parameters, we enable fitting complex personalized models across imaging modalities (fMRI, MEG, EEG) and modeling scales (global, regional, connectivity), while maintaining physiological interpretability essential for clinical translation. Our structured workflow—exploration, optimization, refinement—provides an accessible path through increasingly complex parameter spaces. Key challenges remain where automatic differentiation fails at dynamical bifurcations, precisely where many models operate optimally. Future developments based on shadowing methods or gradient flossing may address these instabilities. Bridging disparate timescales of neural dynamics requires new approaches to parameter decoupling and multi-step optimization. Looking forward, the gradient infrastructure established for optimization enables probabilistic modeling approaches including variational inference and Hamiltonian Monte Carlo [52,53], crucial for clinical applications requiring uncertainty quantification. While computationally demanding, our performance improvements make these increasingly feasible. As computational neuroscience moves toward clinical translation, tools balancing biological realism, computational efficiency, and statistical rigor prove essential. TVB-Optim represents a step toward making personalized brain modeling a practical clinical reality.

## 4 Methods and Materials

### 4.1 Brain Network Models

*Brain Network Model* (BNM), the term used throughout this work, is a broad model category and appears in the literature under different names like *The Virtual Brain* [18], *Connectome-Based Neural Mass Model* [19], *Large-Scale Brain Network Model* [10]. In mathematical terms, a BNM is a system of Stochastic Delay Differential Equation (SDDE), with a formal definition given in equation Equation 4. The equation describes the state of a single node **S**_*i*_ in a network defined by the structural connectivity (cf. Section 4.2.1) matrix *C*. Intrinsic node dynamics *f* are subject to delayed coupling *g* reflecting long-range white matter connections. The whole system is driven by external stimuli *I* and noise *η*.

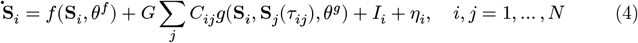

#### 4.1.1 Neural Mass Models

The node dynamics *f* are based on *Neural Mass Models* (NMM), which capture average quantities of populations of spiking neurons [54]. NMMs can be derived in various ways, starting by taking canonical models from dynamical systems literature, like the normal form of the Hopf bifurcation (cf. Section 9.1.3) [55]. Other models are derived by mean-field approaches, employing heuristics, like the Jansen-Rit model [33] (cf. Section 9.1.1) or the reduced Wong Wang model [45] (cf. Section 9.1.2). Modern approaches are able to produce exact reductions of specific spiking models under certain assumptions, like the Montbrió-Pazó-Roxin model for populations of linear integrate and fire neurons [56].

#### 4.1.2 Observation Models

The output of most NMMs is often not directly observable by current imaging techniques, prompting the use of further postprocessing models, here termed *Observation Models*. The observation model **O** can be expressed as a function that transforms the raw state of a BNM into a set of physiological variables of interest *S*_VOI_. These transforms can be layered on top of each other to create multiple complex observations.

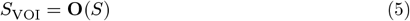

One common transformation used for modelling electromagnetic field measurements like EEG or MEG are leadfield matrices, used to project neural activity from source into sensor space to account for the constraints that the skull imposes on signal measurement. Several methods exist for forward modelling as well as the inverse problem, ranging from mechanistic [57] to Bayesian [58] approaches.

Another target imaging technique is BOLD fMRI that can be obtained from raw neural activity using a Balloon-Windkessel Model [59], acting effectively as a low-pass filter.

#### 4.1.3 Parameter Types - Global, Regional, Structural

BNMs can be parameterized at different levels of granularity, each offering distinct modeling capabilities. Understanding these parameter types is essential for choosing appropriate modeling strategies for specific research questions.

##### Global Parameters

These homogenous parameters are applied consistently across all brain regions, defining the general dynamical regime of the entire system. Examples include global coupling strength between regions, overall neural excitability, or noise levels.

##### Regional Parameters

These heterogeneous parameters allow each brain region to have distinct values, capturing local variations in neural dynamics. For instance, each region might have its own excitability level, time constant, or inhibitory-excitatory balance. This parameterization level acknowledges that different brain areas have specialized functions, resulting in parameter vectors with dimensionality equal to the number of brain regions.

##### Connectivity Parameters

The most granular level involves parameters that directly modify the connectivity structure between regions. Rather than using empirical connectivity weights as fixed constraints, connectivity-scale parameters allow optimization of individual connection strengths. This approach can capture individual differences in brain organization or compensate for measurement noise in structural data or effects of interventions. The parameter space can scale up to the number of possible connections (N^2^ for N regions), though in practice, sparsity constraints or symmetry often reduce this number.

Figure 7 illustrates these three parameterization levels, showing how the same brain network can be described with increasing levels of detail. The choice of parameterization level depends on available data, computational resources, and the specific hypotheses being tested.

**Figure 7.**
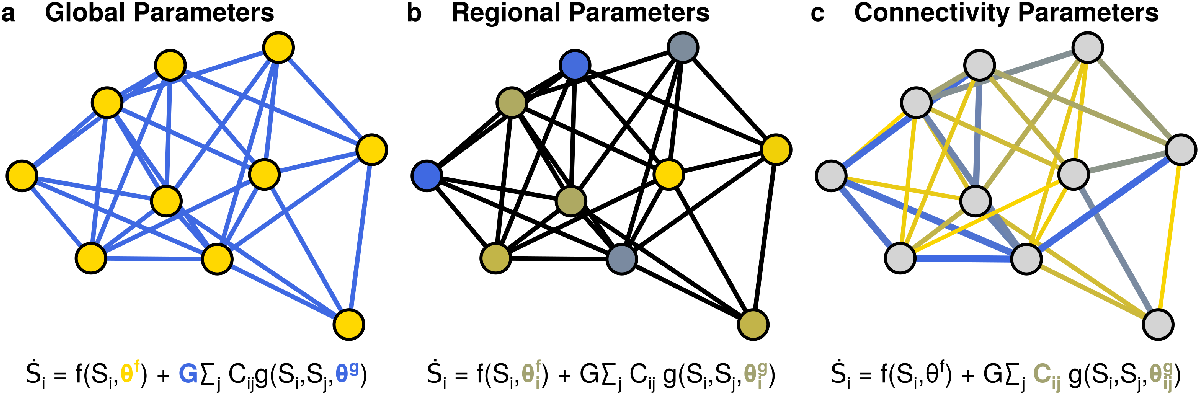
Panel **a**: Global Parameters - uniform parameters applied consistently across all regions of the BNM. Panel **b**: Regional Parameters - heterogeneous, region-specific parameters allowing for spatial variation across different regions of the model. Panel **c**: Connectivity Parameters - directly modify the underlying connectome, affecting the network architecture.

#### 4.1.4 Solving SDDEs

The system of continuous equations defined by Equation 4 can be solved by a basic discretization approach, commonly known as Euler-Maruyama method [60]. Choosing the timestep *dt* directly impacts the accuracy of the solution, with stiff systems requiring smaller timesteps. For properly capturing delay dynamics it should also hold that *dt* is smaller than the smallest delay *τ*. Given a desired simulation time *T*_sim_, this yields a number of steps to be performed depending on the timestep *dt* giving the total computational cost. For better convergence of the deterministic dynamics higher order methods can be used. During this work we use a second-order Runge-Kutta method, also known as Heun’s method, instead of the simpler first-order Euler method. Additionally, we assume that aggregated coupling is slow changing (*g*(*t*) ≈ *g*(*t* + *dt*)) and only evaluated once for each timestep. This is important for performance, as with growing network size *N* the coupling function *g* becomes the dominating factor with node dynamics *f* taking negligible computational time. Delayed states *S*(*t*−*τ*) are supplied via a dedicated history, that has to be provided for each simulation in addition to the state vector as initial condition. The history is an array of length *n*_*τ*_ = *τ*_*max*_/*dt*. Defining the history as simple indexing into an array has the side effect that we can not differentiate with respect to conduction speed defining the time delays via tract length. This is limitation is a tradeoff for performance, which can be lifted switching to a history based on interpolation, also allowing for solvers using adaptive time stepping.

### 4.2 Normative Data

We used the HCP Young Adult dataset (n=650, ages 22-35) acquired on a 3T Siemens Skyra scanner with customized gradients, including resting-state fMRI (2mm isotropic, 0.73s TR), diffusion-weighted MRI (1.25mm isotropic, 128 directions), and structural MRI (0.7mm isotropic T1w/T2w) [61]. Data were preprocessed using standard HCP pipelines and analyzed with the *MRtrix3* toolbox with a detailed description in [38]. The data used for this study was averaged, is purely normative and does not contain any personal information.

#### 4.2.1 Structural Connectivity

From the reconstructed average tractogram, we created a structural connectivity by counting the number of connections between 84 cortical and subcortical regions, that were defined by the Desikan-Killiany atlas [62]. The resulting matrix is shown in Figure 8, panel **a** and was normalized so all weights are in the interval [0, 1]. The average length of the tracts connecting region is shown in panel **b**, which, combined with a conduction speed, set to *v*_*c*_ = 3 m/s by default, can be used to calculate the time delay *τ* between regions.

**Figure 8.**
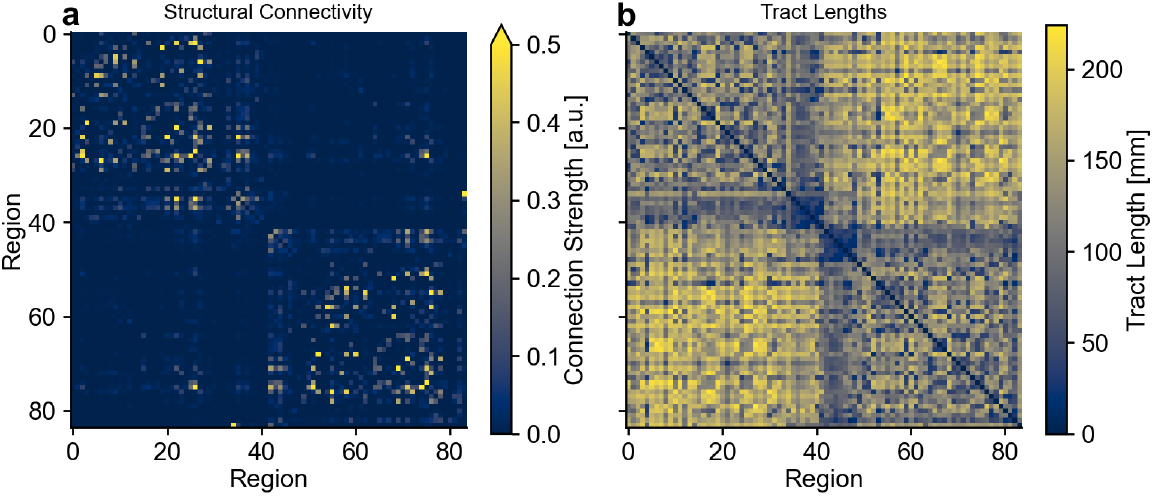
Panel **a** shows the structural connectivity with weights in [0, 1]. For visualization purposes the colormap is clipped at 0.5. Panel **b** shows the tract length in mm.

#### 4.2.2 Functional Connectivity and its Dynamics

FC captures statistical dependencies between spatially remote neurophysiological events [63] and provides a robust and well-studied observable measure in neuroscience [3]. Panel **a** in Figure 9 shows the normative resting state FC used as optimization target throughout this study. Static FC matrices represent time-averaged correlations across entire recording sessions, effectively collapsing temporal information into a single summary statistic.

**Figure 9.**
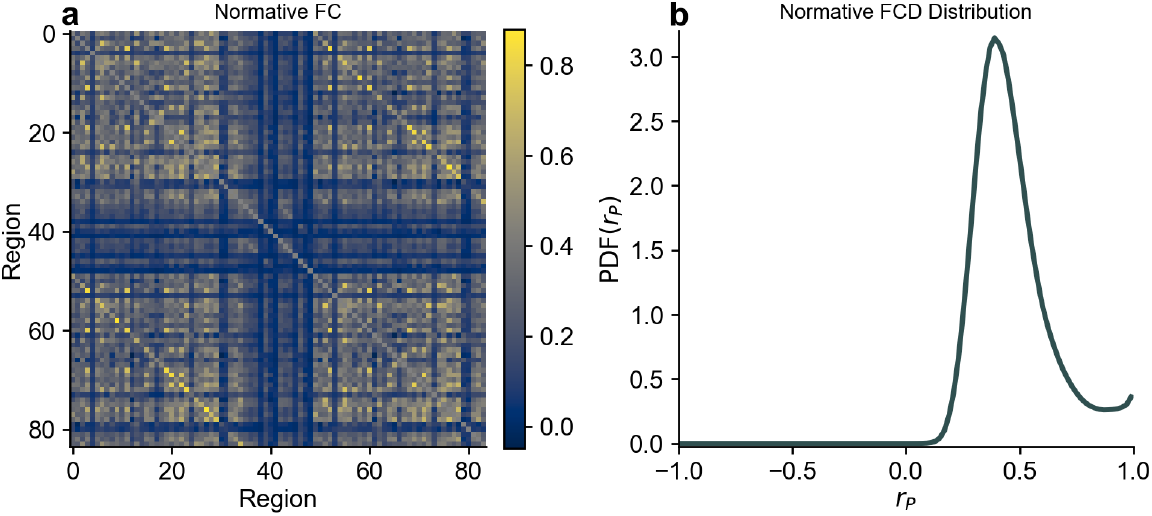
Panel **a** shows the normative FC and panel **b** the normative FCD distribution. Both are parcellated based on the Desikan-Killiany atlas.

This temporal averaging obscures the brain’s dynamic reorganization across different functional states. To quantify these dynamics, we employ FCD [30], which tracks how connectivity patterns evolve over time. The computation proceeds in three steps. First, the time series is divided into overlapping windows (typically 30-60 seconds). Second, a separate FC matrix is computed within each window, yielding a sequence of connectivity snapshots. Third, these FC matrices are vectorized and correlated pairwise, producing the FCD matrix where each entry represents the similarity between connectivity patterns at different time points. High FCD values indicate stable connectivity configurations, while low values reflect transitions between distinct network states. Panel **b** in Figure 9 shows a normative resting state FCD distribution with its characteristic peak around *ρ*_*P*_ ≈ 0.5, indicating balanced dynamics between stability and flexibility, used as target throughout this study.

#### 4.2.3 MEG Peak Frequency Distribution

In resting state, the peak frequency in MEG has been found to negatively correlate with the distance to the visual cortex [32]. We use this finding to create a synthetic target for optimization. In the Desikan-Killiany parcellation, used throughout this study, the lateral occipital gyrus (cf. Figure 10 panel **a** black region) best matches the visual cortex and is assigned the highest peak frequency of *f*_max_ = 11 Hz. The tract length *l* for each region to the lateral occipital gyrus is used to scale down *f*_max_ following equation Equation 6, leading to a minimal peak frequency of *f*_min_ = 7 Hz at the maximum distance *l*_max_. Panel **b** in figure Figure 10 shows the resulting target peak frequency distribution across the cortex.

**Figure 10.**
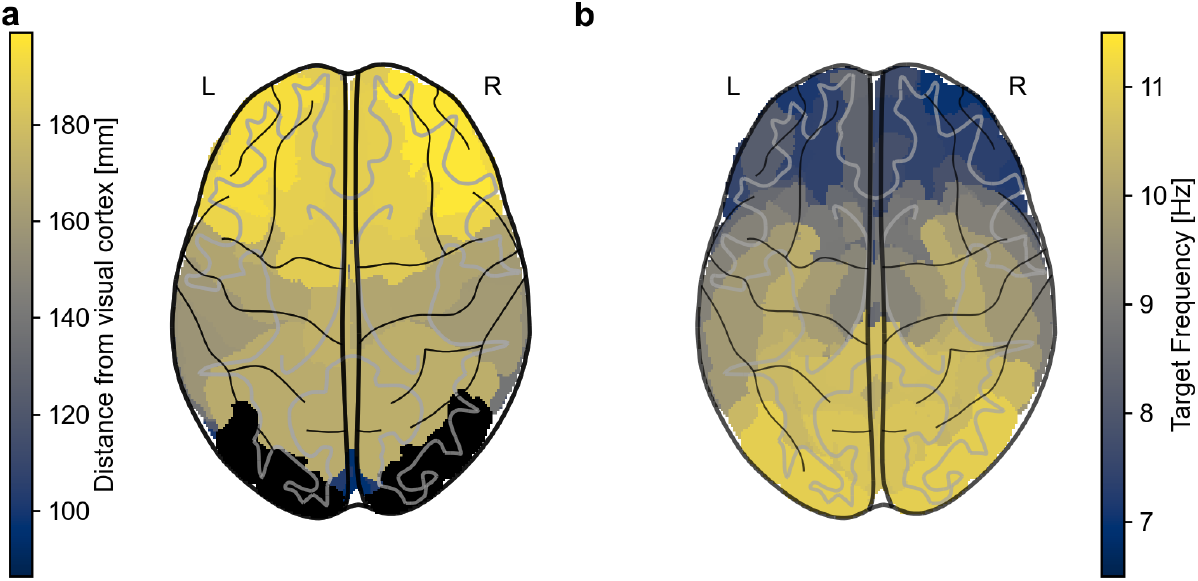
In the Desikian Killian parcellation the lateral occipital gyrus (black) corresponds closest to the visual cortex. All other distance shown in panel **a** are calculated as mean white matter fiber lengths from any region to it. The distance is mapped in panel **b** to a peak frequency with 7Hz for the furthest regions and 11Hz for the closest.

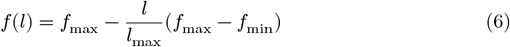

### 4.3 Differentiable Brain Network Models

Efficient parameter estimation requires efficient, fast, numerical simulations, as they accelerate all subsequent estimation methods. While fast simulations provide the foundation, algorithmic choices become the decisive factor in solving estimation problems efficiently. We address this challenge through two approaches: generating optimized code for all models and ensuring their differentiability to enable efficient gradient-based estimation methods.

#### 4.3.1 TVB-O: Easy Model Definition & Fast Code Generation

We use TVB-O [27], leveraging its structured and curated knowledge base for brain simulation, to generate optimized, differentiable code. TVB-O includes a unified interface where the most prominent NMMs (all of Section 9.1) are curated and new ones can be added via a YAML specification. Existing experiments using TVB can be imported via a `.from_tvb_simulator()`function and easily used and extended with new features. New Experiments can be defined directly using TVB-O as shown in Figure 13.

Through templating the structured information can be rendered to simulation code in a language agnostic way. We contributed templates for efficient simulation code, based on the *JAX* library [26], that is completely differentiable via automatic differentiation and ready for use on distributed systems and GPUs. On that basis we build TVB-Optim to provide further utilities to leverage all those capabilities in a user-friendly way. The same unified model specification enables multiple computational workflows, the JAX templates support parameter fitting and optimization, while the Julia templates facilitate bifurcation analysis (Section 9.1.2) via *BifurcationKit*.*jl* [64], demonstrating the versatility of TVB-O‘s language-agnostic approach.

#### 4.3.2 JAX Automatic Differentiation

Automatic differentiation (AD) provides a computationally efficient framework for calculating derivatives of composite functions by methodically applying the chain rule to elementary operations[66]. For composite functions expressed as *f*(*x*) = *f*_*n*_(*f*_*n*−1_(…*f*_1_(*x*))), AD systematically tracks intermediate variables and their corresponding derivatives to determine *∂f*/*∂x* with machine precision.

Unlike numerical methods that approximate derivatives through finite differences, AD computes exact derivatives without introducing truncation or round-off errors. AD also offers substantial advantages over symbolic approaches[16] by processing concrete numerical values rather than manipulating mathematical expressions, thereby efficiently handling complex computational workflows with conditional branches and iterative procedures.

Two principal variants of AD exist: forward mode, which propagates derivatives alongside function values from inputs to outputs; and reverse mode, which first executes a forward computation while recording intermediate values, then propagates gradients backwards from outputs to inputs[65]. The computational efficiency of these approaches varies with problem dimensionality. For functions mapping ℝ^*n*^ to ℝ^*m*^, forward mode requires *n* passes to compute the complete Jacobian, whereas reverse mode necessitates only *m* passes, conferring significant advantages when *m* ≪ *n*[16]. This explains the prevalence of reverse mode in machine learning applications, where the output is typically a scalar loss function (*m* = 1) and the input dimension corresponds to numerous model parameters.

The memory requirements of these approaches can differ substantially because reverse mode incurs additional memory overhead to store intermediate values for the backward pass, while forward mode requires less additional memory. Our implementation leverages JAX’s automatic differentiation capabilities through its gradient and Jacobian-vector product transformations[26], predominantly employing reverse-mode differentiation.

#### 4.3.3 TVB-Optim: A Functional Optimization Framework

The TVB-Optim framework introduces a functional approach to BNM optimization, where all operations are expressed as functions of state, with states represented as JAX PyTrees, a flexible tree-like data structure that can contain nested combinations of arrays, dictionaries, and other Python objects. This design paradigm enables seamless integration with JAX’s automatic differentiation capabilities while maintaining computational efficiency and scalability.

##### Functional State Architecture and Entry Point

The framework’s entry point is the `prepare` function, which takes a TVB-O `SimulationExperiment` (example in Figure 13) and returns a functional simulation interface. Alternatively, models can also be directly defined in TVB-Optim, giving low-level controls over the simulation process making it easy to explore novel model architectures.

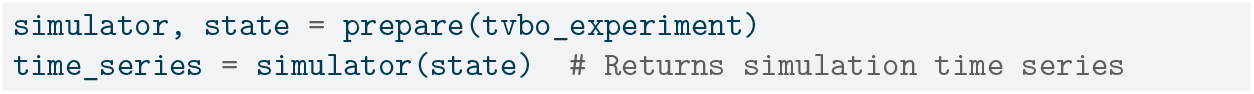

##### Creating Custom Analysis Functions

New analysis functions can be created by wrapping the simulation function. This approach leverages JAX’s familiar NumPy and Scipy interface to compute derivative observations such as power spectra.

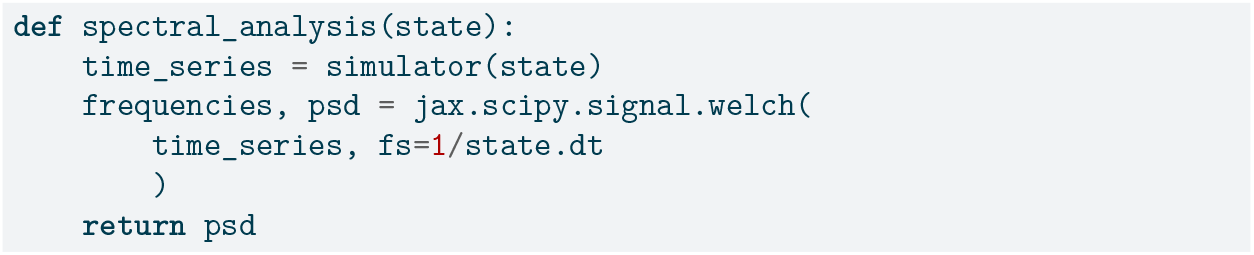

##### Loss Function Definition and Parameter Optimization

Loss functions compare computed derivatives against empirical data. Here we define, in the same way by wrapping the analysis function and enclosing the target data, a spectral correlation loss that measures similarity between simulated and target power spectra.

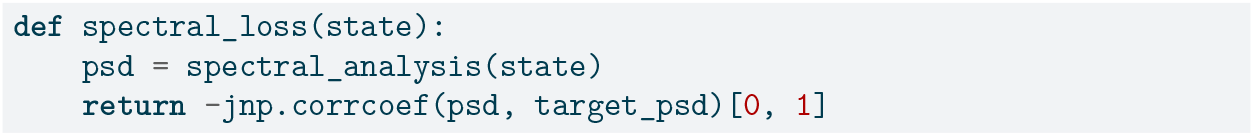

##### Parameter System and Constraints

The so called state contains all model parameters and initial conditions, it supports intuitive dot notation syntax for accessing nested parameters. Array elements within the state can be converted to optimizable parameters. The framework provides specialized parameter types for constraint enforcement.

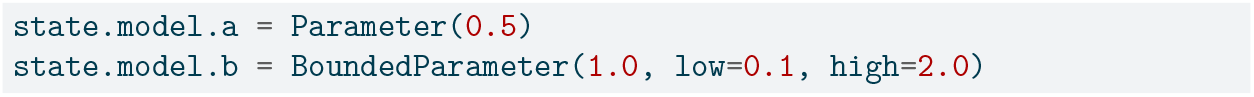

##### Optimization with Optax

Optimizations are performed using the Optax library [67] through the `OptaxOptimizer` class, which supports various gradient-based optimization algorithms. It provides severals convenience functions for debugging and visualizing the optimization process.

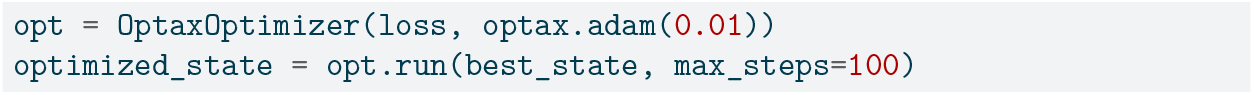

##### Parameter Space Exploration

The framework supports systematic parameter space exploration through an axis system. An axis like `GridAxis` can be placed in the state and `Space` creates an iterable collection of state combinations either as cartesian product or in a zip fashion.

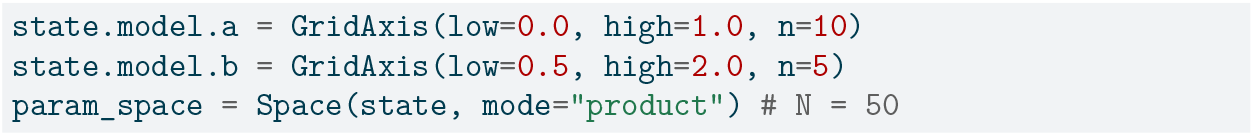

##### Parallel Execution Strategies

TVB-Optim provides execution strategies for parameter space analysis with automatic JAX vectorization for multi-core processing and process mapping for multi-node distribution.

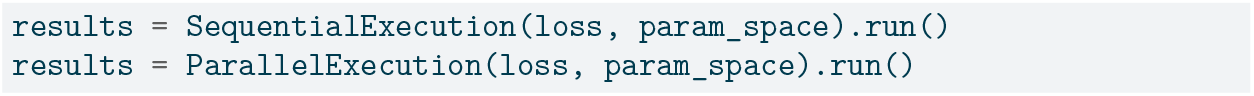

##### Documentation and Reproducibility

All examples and use cases presented in this paper are available as executable documentation in the project’s GitHub repository, providing additional implementation details and extended tutorials for practical application of the framework.

### 4.4 Gradient-Based Optimization

#### 4.4.1 Problem Definition - Loss Function Definition

Parameter optimization is central to personalizing BNMs for clinical applications such as diagnosing or predicting outcomes in various diseases [39]. The fundamental challenge is formalized in equation Equation 7. The loss function ℒ(*θ*) represents a BNM and an observation model **O** which generate predictions based on parameters *θ*. These predictions are then evaluated against empirical observations **Ô** using an appropriate function ℳ to quantify the discrepancy. We seek to identify optimal parameter configurations 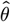 that minimize this loss function across the parameter space.

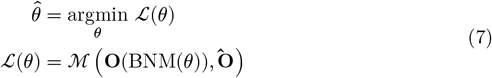

While we frame this as a minimization problem, the approach is equally valid when formulated as maximization. The specific structure of the loss function must be carefully designed to reflect the biological system under investigation. Complex biological phenomena often require composite loss functions, which can be constructed through addition (equation Equation 8) or other differentiable operations, enabling incremental approaches to capture multifaceted biological problems.

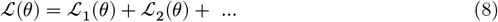

#### 4.4.2 Specific Loss Functions

Having defined the empirical data targets (Section 4.2), we now specify the loss functions used to quantify the match between model predictions and observations. These functions are designed to capture different aspects of brain dynamics across multiple scales and modalities.

##### Functional Connectivity Loss Functions

We employ multiple loss functions to measure different aspects of model performance for FC and FCD.

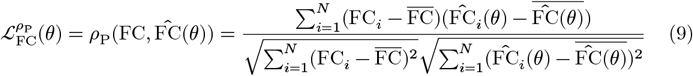

In Equation 9 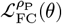 quantifies the Pearson correlation between empirical functional connectivity matrices *F C* and model-predicted matrices 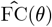. The model parameters are denoted by *θ*, while 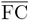 and 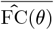 represent the mean values across all *N* regions.

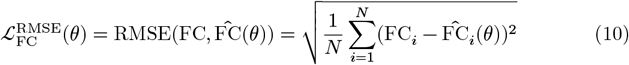

The second loss in Equation 10 measures the root mean square error between empirical and predicted functional connectivity values across all *N* regions, providing a direct measure of prediction accuracy in the original units.

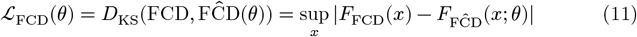

We evaluate the dynamics of FC in Equation 11 using the Kolmogorov-Smirnov statistic *D*_KS_, which computes the maximum distance between the cumulative distribution functions *F*_FCD_(*x*) and *F*_FĈD_ (*x*; *θ*) of the empirical and predicted functional connectivity dynamics, respectively.

##### MEG Spectral Loss Function

To quantify how well our model captures spectral characteristics, we introduce another loss function based on power spectra.

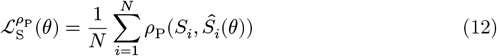

Here, 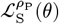 represents the average Pearson correlation between empirical power spectra *S*_*i*_ and model-predicted spectra *Ŝ*_*i*_ (*θ*) across all *N* brain regions. This metric is sensitive to the shape of the power spectrum, putting less emphasis on the magnitude of peaks.

#### 4.4.3 Gradient-Based Optimizers

Gradient-based optimization forms the backbone of modern ML and the methods used throughout this work. Optimizers using the gradient *g*_*i*_ at a certain state *θ*_*i*_ can be written in a general form as Equation 13, where the next state *θ*_*i*+1_ gets updated depending on a potentially adaptive learning rate *α*_*i*_ and the potentially transformed gradient *m*_*i*_(*g*_*i*_).

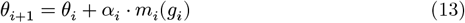

The simplest version is stochastic gradient descent (SGD), where *m*_*i*_(*g*_*i*_) = *g*_*i*_ (no transformation). While effective despite its simplicity [68], SGD can get stuck in local optima, especially for low-dimensional problems.

More sophisticated optimizers improve upon SGD through two key mechanisms: adaptive learning rates and momentum. ADAM [69] combines both by incorporating momentum (first moment estimation) with adaptive learning rates based on gradient magnitudes (second moment estimation), making it popular for its speed and robustness across different optimization landscapes.

Other notable optimizers include RMSProp [70] (adaptive learning rates only), ADAMAX, and ADABelief [71] (belief-based gradient scaling). We used the *Optax* library [67] that provides a broad range of optimizers expressed as flexible gradient transformations that can be easily combined.

#### 4.4.4 Optimization Setting Summary

The optimization approach varied across neural mass models due to their different computational requirements and parameter characteristics. Table 1 summarizes the most important optimization settings used in this study.

**Table 1.**
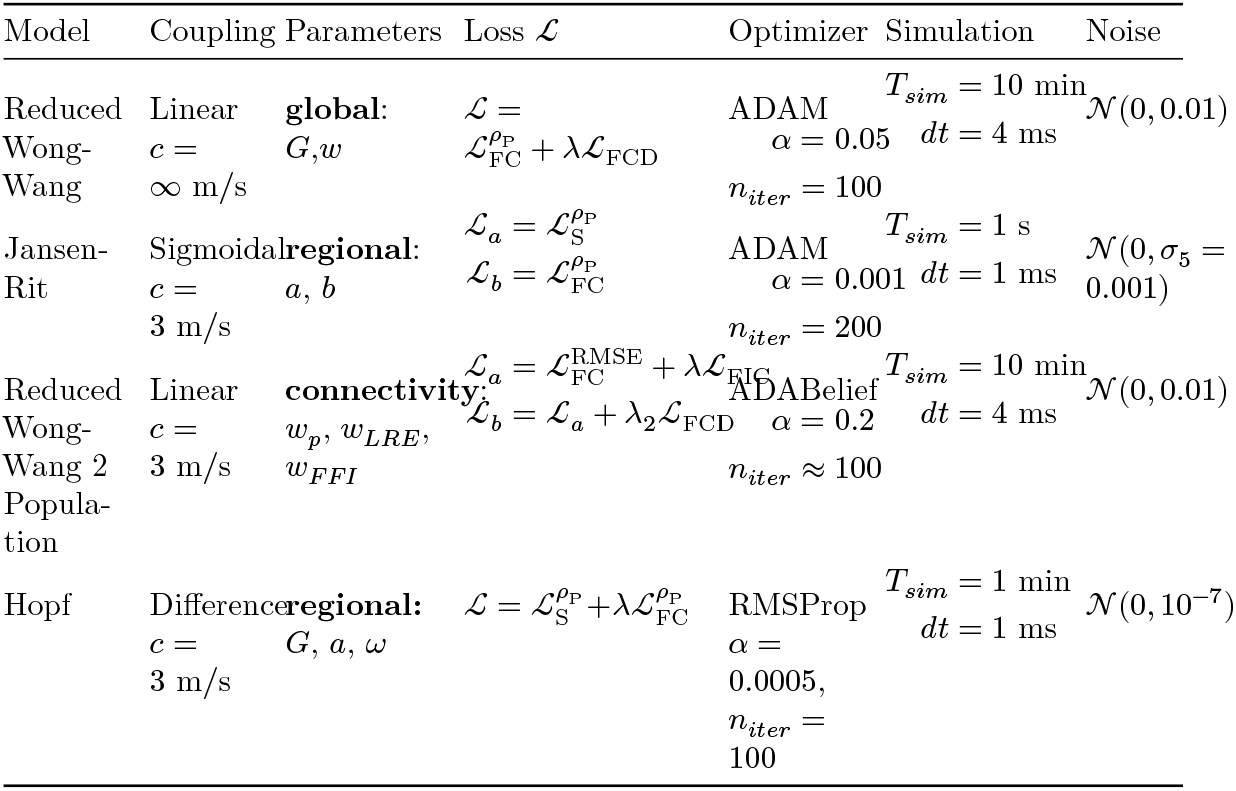
Summary of optimization settings selected for the workflows reported in this paper. Details on models, coupling and meaning of parameters can be found in Section 9.1. A definition of the optimizers can be found in Section 4.4.3.

**Table 2.** Default model parameters for the Jansen-Rit model [33,36]. Parameters marked with _†_ are estimated during the optimization.

| Parameter | Value | Unit | Description |
| --- | --- | --- | --- |
| $A$ | 3.25 | mV | Maximum amplitude of EPSP [mV] |
| $B$ | 22.0 | mV | Maximum amplitude of IPSP [mV] |
| $J$ | 135.0 | [a.u.] | Average number of synapses between three neuronal populations of the model |
| $a_1$ | 1.0 | [a.u.] | Average probability constant of the number of synapses made by the pyramidal cells to the dendrites of the excitatory interneurons (feedback excitatory loop) |
| $a_2$ | 0.8 | [a.u.] | Average probability constant of the number of synapses made by the EINs to the dendrites of the PCs |
| $a_3$ | 0.25 | [*a.u.*] | Average probability constant of the number of synapses made by the PCs to the dendrites of the IINs |
| $a_4$ | 0.25 | [*a.u.*] | Average probability constant of the number of synapses made by the IINs to the dendrites of the PCs |
| $a^\dagger$ | 0.1 | ms<sup>-1</sup> | Reciprocal of the time constant of passive membrane and all other spatially distributed delays in the dendritic network. Also called average synaptic time constant. |
| $b^\dagger$ | 0.05 | ms<sup>-1</sup> | Rate constant of the inhibitory post-synaptic potential (IPSP) |
| $I_{ext}$ | 0.15 | ms<sup>-1</sup> | Mean excitatory external input to the derivative of the state-variable $S_4$ (PCs) represented by a pulse density, that consists of activity originating from adjacent and more distant cortical columns, as well as from subcortical structures |
| $\nu_{max}$ | 0.0025 | ms<sup>-1</sup> | Asymptotic of the sigmoid function<br>Sigm_JR corresponds to the maximum firing rate of the neural populations |
| $v_0$ | 5.52 | mV | Average firing threshold (PSP) for which half of the firing rate is achieved |
| $r$ | 0.56 | mV<sup>-1</sup> | Steepness (or gain) parameter of the sigmoid function Sigm_JR |
| $c_{max}$ | 0.005 | [*a.u.*] | Maximum coupling strength in the sigmoidal coupling function |
| $c_{min}$ | 0.0 | [*a.u.*] | Minimum coupling strength in the sigmoidal coupling function |
| midpoint | 6.0 | mV | Threshold value in the sigmoidal coupling function |

These settings were chosen based on preliminary experiments and computational constraints, with simulation times balanced between accuracy and computational feasibility.

### 4.5 Gradient Analysis and Stability Tools

#### 4.5.1 Gradient Stabilization Techniques

Additional techniques can be used to stabilize the optimization process. One which proved especially useful to deal with large gradients around bifurcation points is gradient clipping [47] where the norm ‖ ⋅ ‖ gradient is clipped to a maximum threshold value (cf. Equation 14).

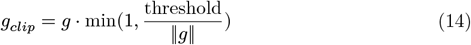

Additionally, the scheduling of the learning rate *α*_*i*_ can be an effective measure to balance of stability by a initially small learning rate and then increasing the learning rate for faster convergence. Decreasing the learning rate after a certain number of iterations can lead to a better final result [72].

#### 4.5.2 Maximum Lyapunov Exponent

The maximum Lyapunov exponent *λ*_*max*_ quantifies the rate of exponential divergence of nearby trajectories in phase space and serves as a key indicator of chaotic behavior in dynamical systems.

We employ the standard algorithm [73] for computing the maximum Lyapunov exponent by evolving the system in parallel with infinitesimally displaced initial conditions. Starting with the original trajectory **S**(*t*) and a nearby trajectory **S**(*t*) + *δ***S**(*t*) where |*δ***S**(0)| = *ϵ*_0_ is small, we integrate both trajectories forward in time. After a fixed number of integration steps, we measure the separation |*δ***S**(*t*)| between the trajectories. To prevent numerical overflow and maintain the linear approximation, we rescale the displacement vector by setting *δ***S**(*t*) ← *δ***S**(*t*) ⋅ *ϵ*_0_/|*δ***S**(*t*)|.

The maximum Lyapunov exponent is then calculated as Equation 15, where *N*_*λ*_ is the total number of rescaling events, *T* is the simulation time between rescalings, and *ϵ*_0_ is the initial separation distance. This procedure ensures that we track the average exponential growth rate of small perturbations while avoiding numerical instabilities.

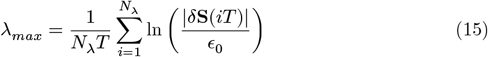

#### 4.5.3 Bifurcation Analysis

Bifurcation analysis serves as a valuable guide for optimization by revealing the qualitative behavior of dynamical systems as parameters change. While computationally limited to 1-2 parameters of a single region, this analysis provides crucial insights into system stability and parameter sensitivity that inform optimization strategies.

##### Single Region Analysis

For detailed bifurcation analysis of individual brain regions, we leverage the computational pipeline from TVB-O to Julia’s BifurcationKit.jl [64]. This approach enables rigorous mathematical analysis of bifurcation points, within single-node dynamics.

##### Network-Level Adiabatic Scanning

At the network level, we employ adiabatic parameter scanning to identify bifurcation phenomena across the full system. This method involves slowly increasing and then decreasing a target parameter while continuously running the simulation, tracking key statistics including mean, minimum, and maximum values of relevant observables. This adiabatic approach helps identify transitions between fixed points and limit cycles, while also revealing phenomena such as hysteresis where the system’s behavior depends on the direction of parameter change. Such insights are invaluable for understanding the dynamic landscape.

#### 4.5.4 Performance Benchmark Design

We conducted comprehensive performance benchmarks to evaluate TVB-Optim across multiple computational dimensions relevant to optimization workflows. All experiments were repeated multiple times to obtain robust statistical estimates.

##### Hardware specifications

CPU benchmarks used a 12th Generation Intel Core i5-1240P mobile processor (4 performance cores, 8 efficiency cores, 16 threads) with 64GB RAM. GPU benchmarks utilized a compute node with 4 Nvidia V100 GPUs (32GB RAM each). These benchmarks demonstrate algorithmic scaling properties and qualitative performance improvements, with absolute values dependent on specific hardware configurations.

##### Forward simulation benchmarks

We compared our JAX implementation against the established TVB framework by simulating all 21 default TVB neural mass models. Each model was run for 100,000 timesteps on 84-region networks with stochastic noise. We tested both with and without axonal transmission delays to assess the impact of history management on performance. Runtime ratios were calculated as TVB execution time divided by JAX execution time.

##### Parallelization benchmarks

We measured scaling behavior by executing *N* = 1024 simulations with varying degrees of parallelization *n*, ranging from sequential execution (*n* = 1) to maximum parallelization (*n* = 128, limited by device memory). Tests covered CPU and single-GPU execution at both 32-bit and 64-bit precision, plus distribution across 4 GPUs to assess multi-device scaling efficiency.

##### Automatic differentiation benchmarks

We evaluated gradient computation efficiency by measuring time and memory usage for forward-mode and reverse-mode AD as functions of parameter count *N*_*θ*_. Tests covered parameter counts from 1 to over 14,000, corresponding to the different parameterization scales demonstrated in Results. Computation times were normalized to single forward simulation cost for comparison.

## 5 Acknowledgements

We acknowledge support by the EU Horizon Europe program Horizon EBRAINS2.0 (101147319), Virtual Brain Twin (101137289), EBRAINS-PREP (101079717), AISN (101057655), EBRAIN-Health (101058516), EIC grant PHRASE (101058240), the Digital Europe Programme TEF-Health (101100700), Shaiped (101195135), CoordinaTEF (101168074); the German Research Foundation SFB 1436 (project ID 425899996), SFB 1315 (project ID 327654276), SFB 936 (project ID 178316478), SFB-TRR 295 (project ID 424778381), SPP Computational Connectomics RI 2073/6-1, RI 2073/10-2, RI 2073/9-1, DFG Clinical Research Group BECAUSE-Y (504745852); Berlin University Alliance OpenMake; the Virtual Research Environment at the Charité Berlin; EBRAINS Health Data Cloud; and the Berlin Institute of Health and Foundation Charité.

## 7 Author Contributions

**M.P**. conceptualization, methodology, software (TVB-Optim), validation, formal analysis, investigation, data curation, writing-original draft, writing-review and editing, visualization. **L.M**. methodology, software (TVB-O), writing-review and editing. **E.R**. conceptualization, methodology, writing-review and editing. **D.P**. conceptualization, supervision, writing-review and editing. **M.S**. conceptualization, data curation, supervision, writing-review and editing. **P.R**. supervision, funding acquisition, writing-review and editing. During the preparation of this work, the authors used Claude Sonnet 4.5 (Anthropic) to improve language and readability. After using this tool, the authors reviewed and edited the content as needed and take full responsibility for the content of the publication.

## 8 Code and Data Availability

The TVB-Optim software is available as an open-source Python library at https://github.com/virtual-twin/tvboptim under the EUPL v1.2 license. All analysis code used to generate the results presented in this manuscript is included in the repository and documented at https://virtual-twin.github.io/tvboptim. Population-averaged empirical neuroimaging data used for model validation are included in the repository. As these data represent population averages rather than individual measurements, they are not subject to data protection regulations.

## 9 Supplementary Materials

### 9.1 Specific Neural Mass Models

Different NMMs are used to demonstrate the capabilities of differentiable BNMs. In the following the exact formulation and parameter values used for each model will be shown.

#### 9.1.1 Jansen-Rit

The Jansen-Rit model [33] describes the dynamic interactions between three populations of cortical neurons within a single cortical column, namely pyramidal cells, excitatory interneurons, and inhibitory interneurons. The model is described by a six-dimensional nonlinear dynamical system parametrized as in [36].

Internal dynamics:

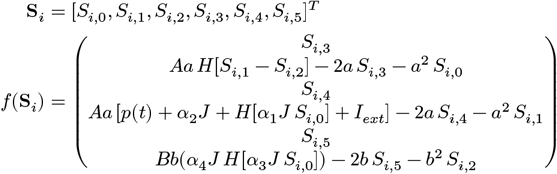

with:

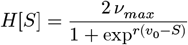

Coupling function:

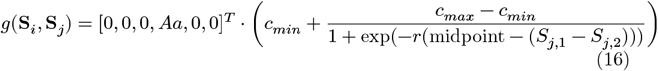

#### 9.1.2 Reduced Wong-Wang

Two different variants of the reduced Wong-Wang model [9,45] are used in this study. A single population version and a two population version with excitatory and inhibitory populations.

##### Single Population

For the single population model, the formulation (Equation 17) and parameters (Table 3) from [30] are used.

**Table 3.** Default model parameters for the reduced Wong-Wang model. Parameters marked with _†_ are estimated during the optimization.

| Parameter | Value | Unit | Description |
| --- | --- | --- | --- |
| $I_{\text{ext}}$ | 0.3 | nA | External input current to the neurons population |
| $J_N$ | 0.2609 | nA | Excitatory recurrence |
| $a$ | 0.27 | pC<sup>-1</sup> | Slope (or gain) parameter of the sigmoid input-output function H |
| $b$ | 0.108 | kHz | Shift parameter of the sigmoid input-output function H |
| $d$ | 154.0 | ms | Scaling parameter of the sigmoid input-output function H |
| $\gamma$ | 0.641 | dimensionless | Kinetic parameter |
| $\tau_s$ | 100.0 | ms | Kinetic parameter |
| $w^\dagger$ | 0.6 | dimensionless | Excitatory recurrence |

Internal dynamics :

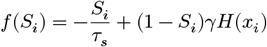

where:

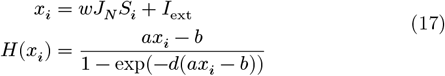

Coupling function :

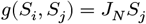

##### Bifurcation Analysis

A bifurcation analysis was performed on model parameters *w* and *I*_ext_, revealing a cusp bifurcation that gives rise to bistability in the model (cf. Figure 12 panel **a** and **b**). This behavior changes when coupled to other populations. For a networked model, standard bifurcation analysis becomes unfeasible, but network behavior can be investigated by performing an adiabatic scan. By slowly increasing and decreasing the global coupling *G* during simulation, the network activity can be observed to switch from the low activity state to the high activity state and back (cf. Figure 12 panel **c** and **d**). Additionally, bistability emerges due to network effects even for settings of *w* that previously did not exhibit bistability.

**Figure 11.**
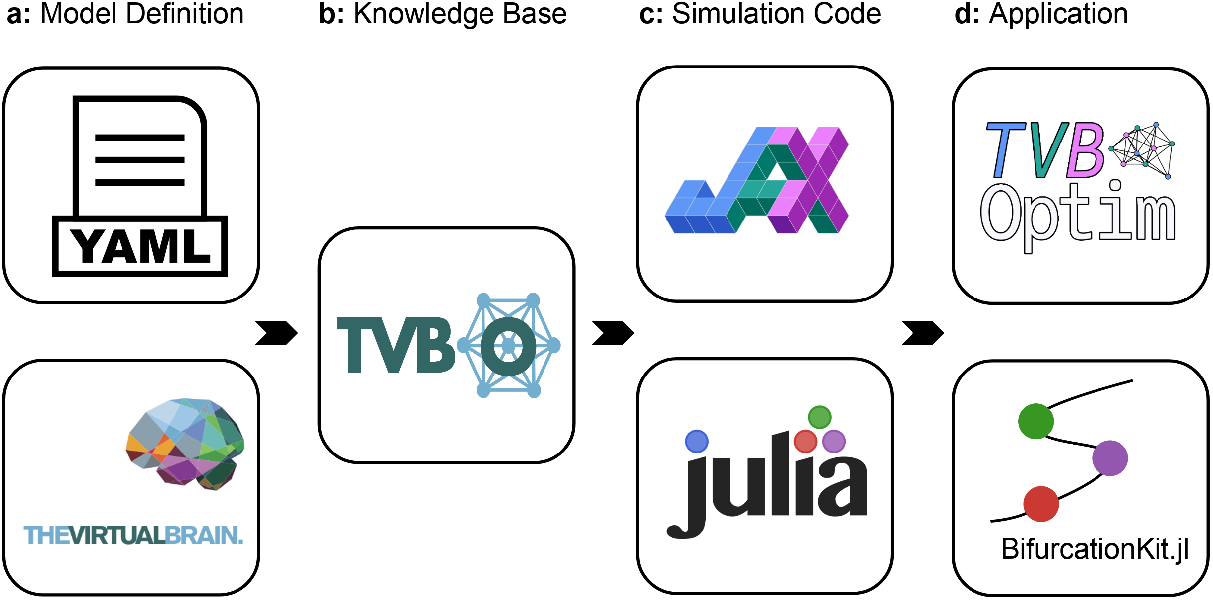
The TVB-O workflow: Panel **a** shows two approaches for TVB-O model definition: using YAML files where novel models can easily be added, or importing from existing TVB simulators to extend current experiments. Panel **b** illustrates how TVB-O‘s structured, semantic BNM representation enables flexible code generation across different target languages such as JAX or Julia (panel **c**). Panel **d** demonstrates downstream applications that utilize this generated code, including TVB-Optim for efficient parameter estimation and BifurcationKit.jl for single-node bifurcation analysis.

**Figure 12.**
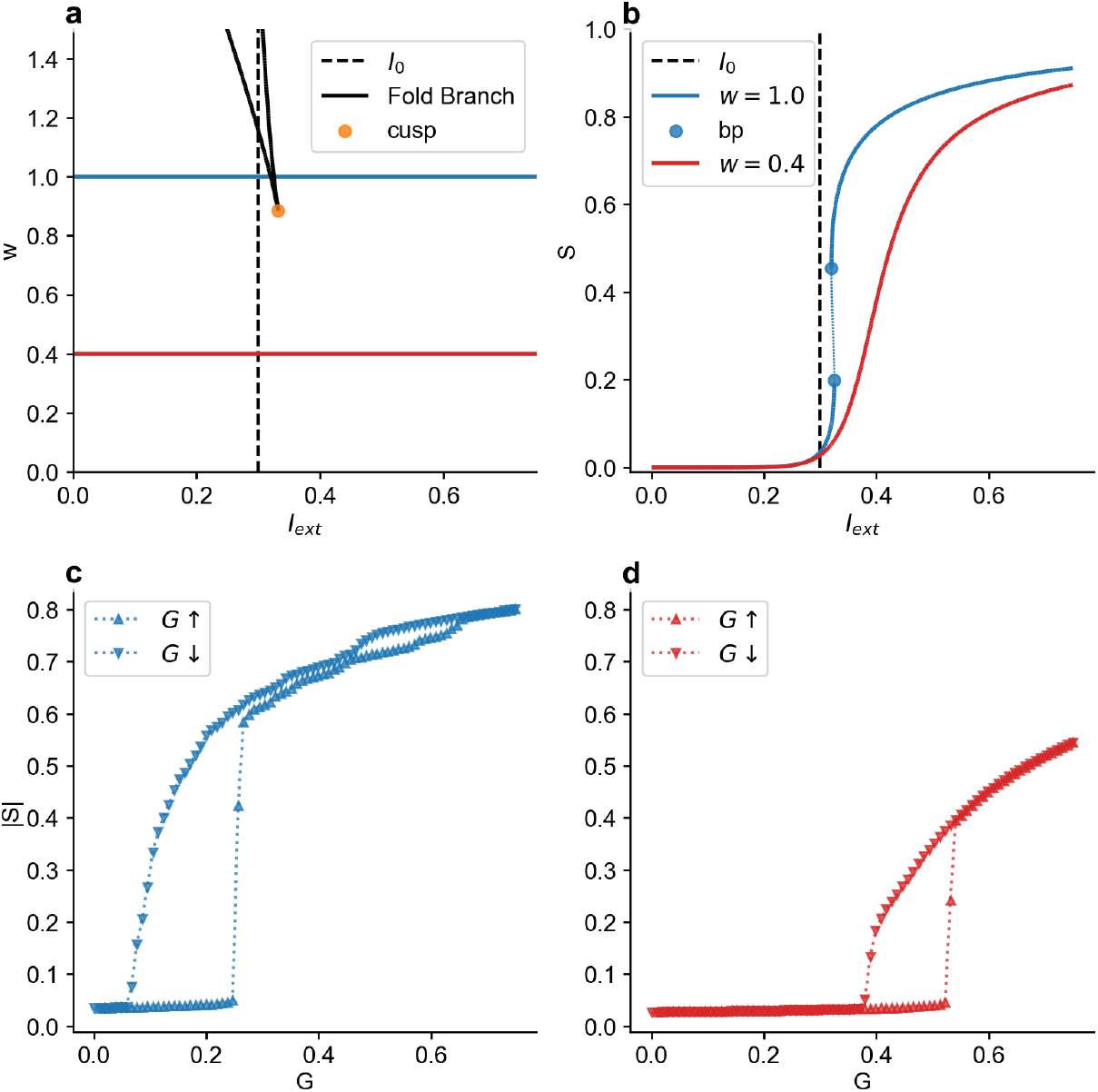
**Top Row:** Bifurcation diagram for the reduced Wong-Wang model. Panel **a** shows a codimension 2 analysis for parameters *w* and *I*_*ext*_ with a cusp bifurcation giving rise to two fold bifurcation branches. Panel **b** shows two cuts through the 2D bifurcation diagram, one at *w* = 1.0 above the cusp bifurcation with two fold bifurcations (bp) and one at *w* = 0.2 below the cusp bifurcation without a bifurcation. **Bottom Row:** Adiabatic scan for the networked reduced Wong-Wang model with *w* = 1.0 (panel **c**) and *w* = 0.2 (panel **d**). Global coupling *G* was slowly increased from 0 to 0.75 and then decreased to 0. Each step was initialized with the state of the previous step. Both panels show the average network activity over the last 1000 time steps. Both scans present a bistability with hysteresis.

##### Two Population Model

For the formulation of the model containing an excitatory and an inhibitory population, the formulation (Equation 18) and parameters (Table 4) from [38] are used.

**Table 4.** Default model parameters for the 2 population reduced Wong-Wang model. Parameters marked with _†_ are estimated during the optimization.

| Parameter | Value | Unit | Description |
| --- | --- | --- | --- |
| $G$ | 2.0 | [a.u.] | Global coupling scaling |
| $I_{ext}$ | 0.0 | [a.u.] | External stimulation parameter for evoked potential simulated activity applied to the excitatory population firing rate IE |
| $I_o$ | 0.382 | nA | Effective external input parameter to the excitatory E and inhibitory I population |
| $J_N$ | 0.15 | [a.u.] | Excitatory synaptic coupling parameter |
| $J_I$ | 1.0 | [a.u.] | Local feedback inhibitory synaptic coupling to the excitatory population firing rate IE |
| $W_E$ | 1.0 | [a.u.] | Excitatory population external input scaling weight |
| $W_I$ | 0.7 | [a.u.] | Inhibitory population external input scaling weight |
| $a_E$ | 310.0 | nC<sup>-1</sup> | Slope (or gain) parameter of the sigmoid function $H_E$ |
| $a_I$ | 615.0 | nC<sup>-1</sup> | Slope (or gain) parameter of the sigmoid function $H_I$ of the inhibitory population |
| $b_E$ | 125.0 | Hz | Shift parameter of the sigmoid function $H_E$ of the excitatory population |
| $b_I$ | 177.0 | Hz | Shift parameter of the sigmoid function $H_I$ of the inhibitory population |
| $d_E$ | 0.16 | s | Scaling parameter of the sigmoid function $H_E$ of the excitatory population |
| $d_I$ | 0.087 | s | Scaling parameter of the sigmoid function $H_I$ of the inhibitory population |
| $\gamma_E$ | 0.000641 | [a.u.] | Excitatory population kinetic parameter |
| $\gamma_I$ | 0.001 | [a.u.] | Inhibitory population kinetic parameter |
| $\lambda$ | 0.0 | [a.u.] | Inhibitory global coupling scaling |
| $\tau_E$ | 100.0 | ms | Kinetic parameter that represents the decay times for NMDA synapses |
| $\tau_I$ | 10.0 | ms | Kinetic parameter that represents the decay times for inhibitory GABA synapses |
| $w_p$ | 1.4 | [a.u.] | Excitatory population recurrence weight |

**Table 5.** Default model parameters for the 2 population reduced Wong-Wang model. Parameters marked with _†_ are estimated during the optimization.

| Parameter | Value | Unit | Description |
| --- | --- | --- | --- |
| $a^\dagger$ | -0.5 | [a.u.] | Local bifurcation parameter |
| $\omega^\dagger$ | 1.0 | [a.u.] | Angular frequency |

Internal dynamics :

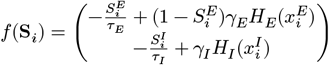

with:

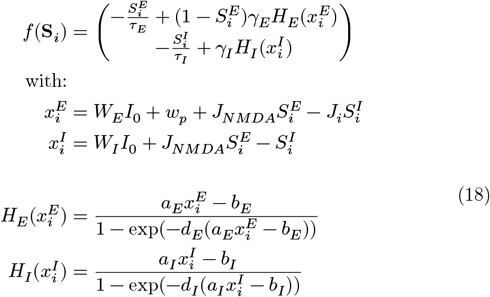

Coupling function :

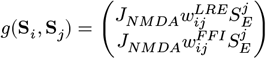

#### 9.1.3 Hopf Normal Form

The Hopf normal form is used as NMM following [40].

Internal dynamics :

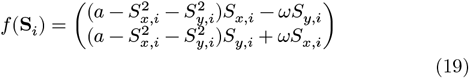

Coupling function:

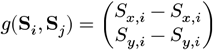

### 9.2 Code Example

The following shows the code for the first result use-case Section 2.1 to get a first impression on how a TVB-Optim workflow looks like. For more information we refer to the documentation, which also contains all other use-cases.

#### 9.2.1 Optimization Workflow

Figure 13 shows the workflow of Figure 1 examplified for the reduced Wong-Wang model to fit FC and FCD data. It was used to create panel **a** and **b** of Figure 2 and performs an optimization starting at the best parameter values from the exploration phase.

**Figure 13.**
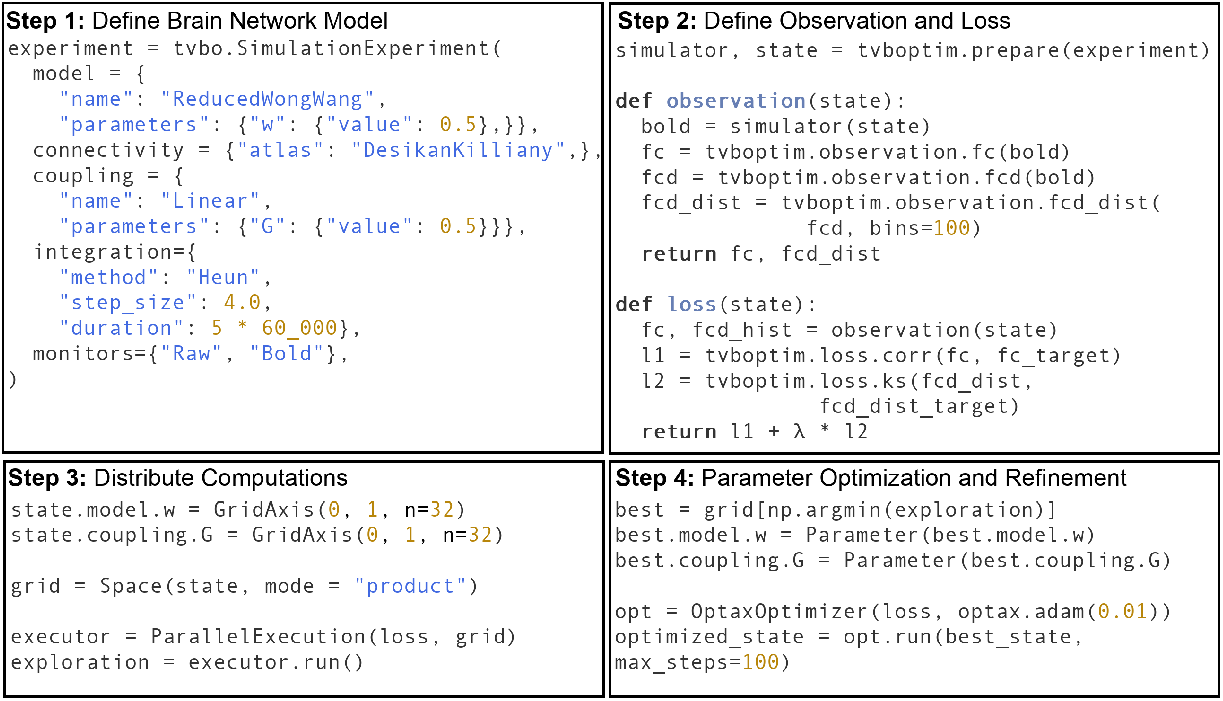
Using a high level description and drawing additional information from the ontology the Reduced Wong-Wang model is placed at each node of the network giving a BNM (panel **1**.). The TVB-O *SimulationExperiment* is transformed into a functional form based of a differentiable *simulator* function and a *state* that contains the parameters, initial conditions and additional information. Building on the *simulator* function, additional functions are defined for observation and loss that in this case compare the FC and FCD histogram to empirical data (panel **2**.). *G* and *w* are set up as GridAxis with 32 elements each (1024 total) participating in the exploration of the grid Space (panel **3**.). For the subsequent optimization, the best values from the exploration are selected and converted into Parameter objects so they are differentiated and updated at each optimization step (panel **4**.).

#### 9.2.2 Parameter Refinement

Some problems require heterogenous parameters, so that different brain regions have different dynamics. This can be achieved by setting the shape from a scalar to a vector of length equal to the number of regions in the BNM, as shown in Figure 14.

**Figure 14.**
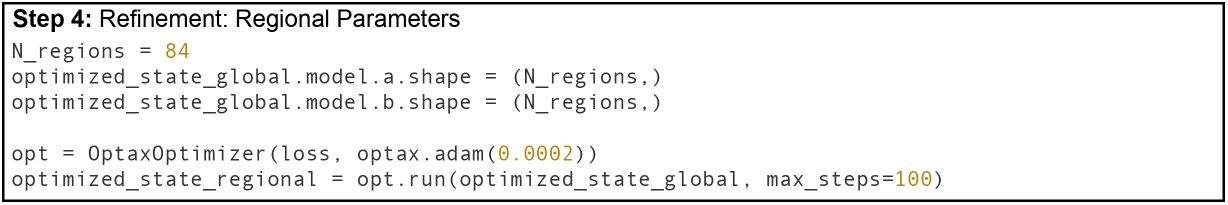
Setting the shape of optimized global parameters to match the number of regions in the BNM turns them into regional parameters. The subsequent optimization can refine the initial fit by exploiting heterogeneity.

### 9.3 Additional Performance Benchmarks

We performed additional benchmarks to highlight interesting and relevant computational behaviors that provide deeper insights into the performance characteristics and tradeoffs chosen for the JAX backend.

#### 9.3.1 Network Size Comparison

To evaluate scalability, we benchmarked runtime performance across different network sizes ranging from N=10 to N=1000 nodes, using a fixed number simulation steps with noise and delay included. The results reveal distinct computational scaling behaviors between the backends.

For small networks, TVB’s Python overhead dominates the runtime, while JAX’s just-in-time compilation eliminates this overhead penalty. JAX runtime grows approximately as *O*(*N* ^2^) with network size, as expected for network simulations, appearing as a straight line on the log-log plot (cf. Figure 15 panel **a**). In contrast, TVB runtime shows minimal growth for smaller networks until *N* ≈ 50, appearing nearly horizontal on the log scale, after which the *O*(*N* ^2^) computational complexity begins to dominate over Python overhead (cf. Figure 15 panel **b**).

**Figure 15.**
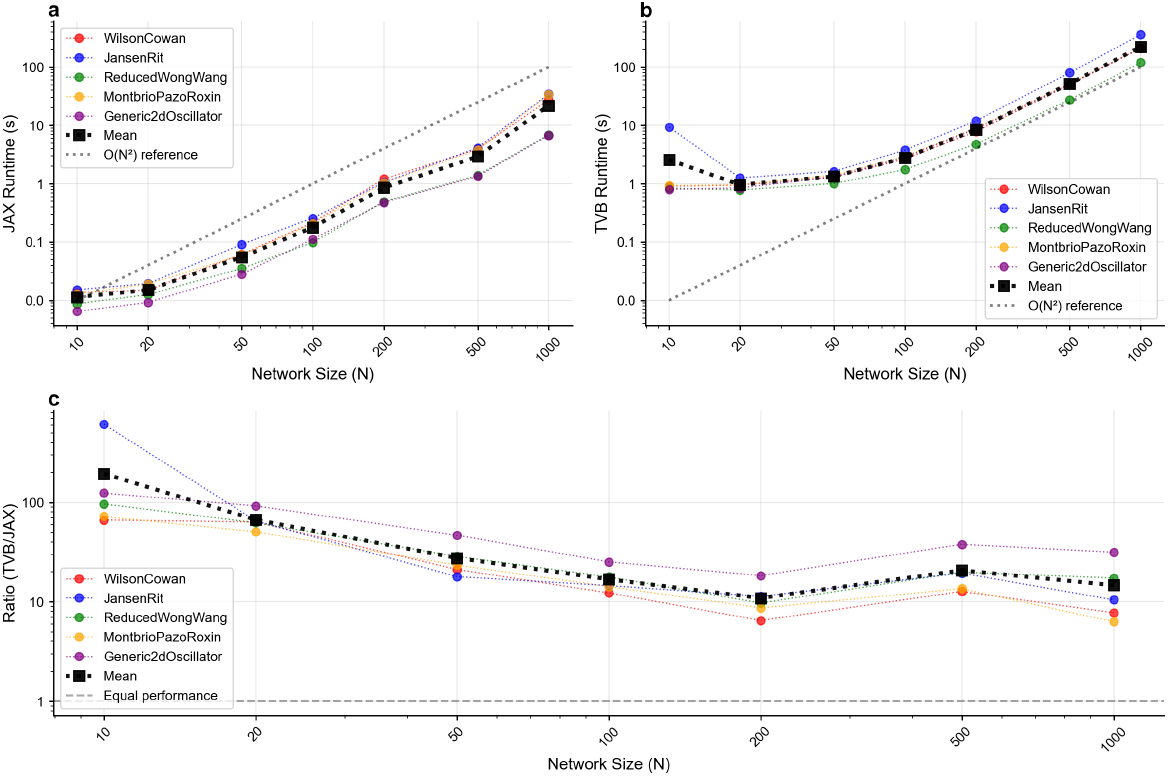
Runtime performance comparison across network sizes. Panel **a** shows JAX backend runtime as a function of network size, scaling approximately as *O*(*N* ^2^) and closely following the theoretical reference line on the log-log plot, indicating efficient computational scaling. Panel **b** displays TVB backend runtime versus network size, showing minimal growth until *N* ≈ 50 on the log scale, suggesting that Python overhead dominates for smaller networks before *O*(*N* ^2^) complexity takes over. Panel **c** presents the runtime ratio (TVB/JAX) across network sizes, demonstrating that JAX maintains consistent performance advantages with over 10× speedup around N=100 and beyond. All simulations used a fixed duration with identical connectivity matrices scaled to different network sizes. Individual model lines show performance for specific neural mass models, while the mean line represents average performance across all models. The O(N^2^) reference line illustrates theoretical quadratic scaling expected for network simulations.

The runtime ratio stabilizes around *N* ≈ 100, where JAX consistently maintains over 10× speedup compared to TVB (cf. Figure 15 panel **c**). Interestingly, for very large networks (N>5000, not shown), the performance difference between simulations with and without delay diminishes in JAX, as the matrix multiplication operations *O*(*N* ^3^) begin to dominate over the history allocation penalty.

#### 9.3.2 Integration Step Size Comparison

We evaluated performance across different integration step sizes dt to understand the computational trade-offs between temporal resolution and runtime efficiency. Beyond the inherent performance benefit that larger step sizes provide by requiring fewer steps to reach the simulation duration T, dt should ideally be chosen as large as numerical stability allows. The benchmarks reveal different behaviors between the two backends across the tested range of step sizes.

TVB maintains mostly constant time per integration step across all step sizes (cf. Figure 16 panel **b**), indicating that its implementation is largely insensitive to temporal resolution changes. In contrast, JAX exhibits significantly slower performance for smaller step sizes (dt < 0.1), but maintains consistent and superior performance for larger step sizes (dt 0.1) (cf. Figure 16 panel **a**).

**Figure 16.**
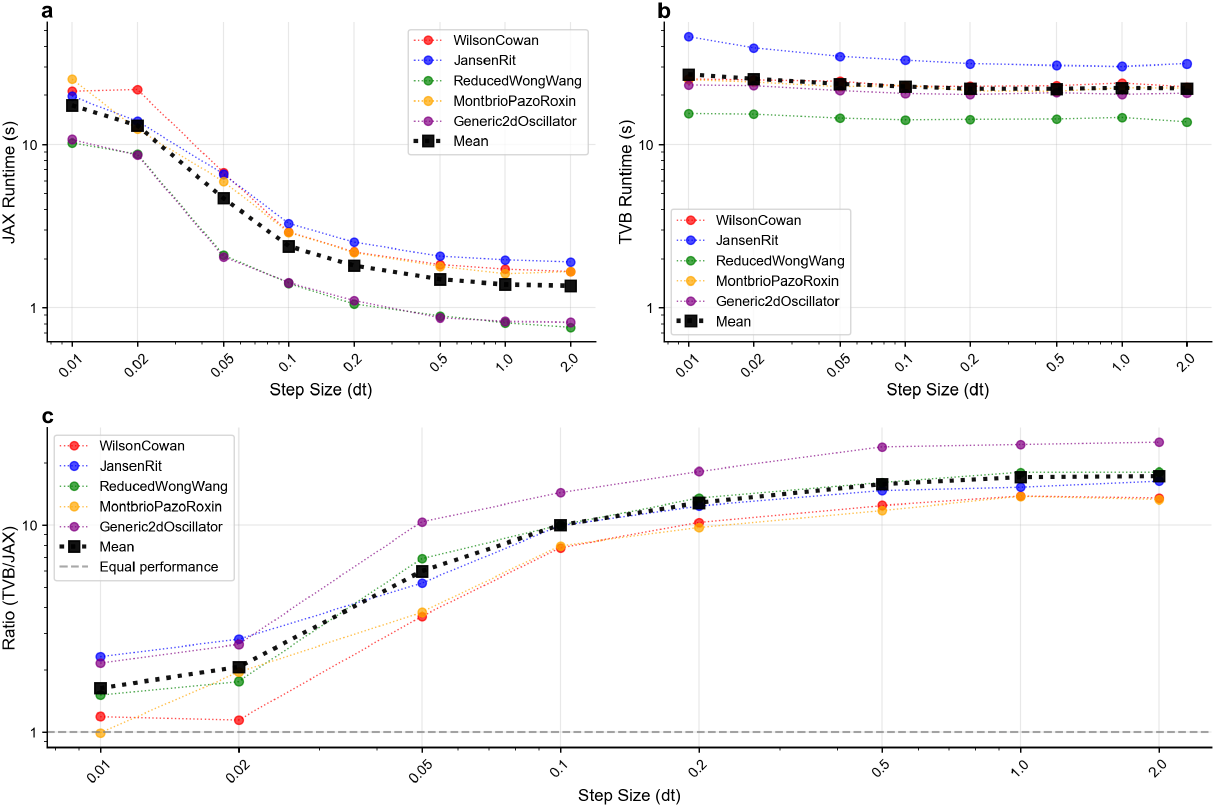
Figure. Runtime performance comparison across integration step sizes. Panel **a** shows JAX backend runtime as a function of integration step size, revealing poor performance for small step sizes (dt < 0.1) but consistent performance for larger step sizes (dt 0.1). Panel **b** displays TVB backend runtime versus step size, showing relatively consistent performance across all temporal resolutions. Panel **c** presents the runtime ratio (TVB/JAX) across step sizes, illustrating that JAX’s performance advantage diminishes significantly for small step sizes where it only marginally outperforms TVB, while maintaining substantial speedups for larger step sizes. All simulations used a fixed number of integration steps (100,000) with a network size of 84 nodes. Individual model lines show performance for specific neural mass models, while the mean line represents average performance across all models. Ratios are calculated from mean runtimes across multiple simulation runs.

This JAX behavior represents a conscious design trade-off optimized for reasonable performance during both forward simulation and backpropagation. Our implementation balances memory usage and computational efficiency, as alternative approaches tend to suffer from explosive RAM requirements for small step sizes combined with longer simulations. To address this limitation, we offer a performance optimization switch (small_dt = True) that resolves the performance degradation for small step sizes, though this option works efficiently only with forward differentiation.

#### 9.3.3 Fundamental performance differences based on coupling type

The coupling type significantly influences the computational complexity of brain network models, as clearly demonstrated in the runtime performance comparisons (cf. Figure 6 panel **a** and **c**). Coupling with time delay (Equation 20) represents the most biologically realistic approach but also the most computationally expensive implementation.

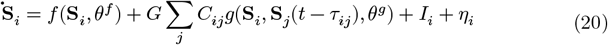

Time-delayed coupling requires individual delay handling for each connection, necessitating gather operations to collect the correct values from a history buffer. This results in **S**_*j*_ being a matrix ∈ ℝ^*N*×*N*^ where *N* defines the number of regions. The coupling function *g* must then be applied to each matrix element, followed by weighting through the connection strength *C*_*ij*_ via element-wise multiplication.

In contrast, coupling without delay (Equation 21) offers computational advantages where **S**_*i*_ and **S**_*j*_ can be vectors ∈ ℝ^*N*^ to which the coupling function *g* is applied, weights can be applied with efficient matrix multiplication.

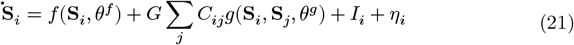

Special cases exist where coupling functions create computational complexity similar to delayed coupling. The most common example is difference coupling (Equation 22), where the interaction between **S**_*i*_ and **S**_*j*_ requires the latter to be a matrix.

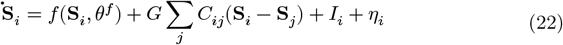

Notably, difference coupling can be mathematically rewritten as linear coupling where the connection weights *C*_*ij*_ are transformed to the graph Laplacian, offering computational optimizations in specific implementations.

#### 9.3.4 Comparison with other frameworks

To establish an upper bound for simulation performance and contextualize our general-purpose JAX implementation, we benchmarked against fastTVB [21], a highly specialized C implementation hand-crafted for a single model configuration. This comparison illustrates the performance ceiling achievable through model-specific optimization versus the trade-offs inherent in maintaining generalizability across neural mass models.

The benchmark employed a simulation configuration using the Reduced Wong-Wang model (cf. Section 9.1) with dt = 0.1 ms, T = 10s duration, 379 brain regions, including both stochastic noise and time-delayed coupling, with BOLD monitor sampling at 720 ms intervals. The performance comparison (Table 6) shows fastTVB achieves the best performance, while our JAX implementation is slower but still substantially faster than the original TVB framework.

**Table 6.** Performance comparison across brain network modeling frameworks for a standardized Reduced Wong-Wang simulation (379 regions, 10s duration, dt=0.1ms, BOLD).

| Framework | Time [s] |
| --- | --- |
| TVB | 208 |
| JAX | 38 |
| fastTVB | 6 |

### 9.4 Landscape of Brain Network Modeling Tools

Several tools have been developed to simulate BNMs. They differ in their target language as well as in their degree of specialization and way to define and simulate BNMs. Loosely, they can be grouped into the following categories:

#### 9.4.1 All-in-one Toolboxes

Tools in this category provide a unified interface to a wide range of different NMMs and postprocessing pipelines. They offer much utility and are well documented for scientific tools.

A pioneering tool in this category is *The Virtual Brain* (TVB) [10,18], which is a *Python* toolbox that has played a crucial role in the development of the field. It sparked a broad ecosystem of adjacent tools and services, from graphical user interfaces to cloud services [75].

Complex tools like TVB have to make opinionated choices regarding the interface and software stack. This led to similar tools, targeting to manage these tradeoffs differently or utilize more modern software stacks. Also in *Python, neurolib* focuses on being more hackable and extendable with included evolutionary optimization algorithms [76]. The *PyRates* project is another tool, taking a graph-based approach with its own domain-specific language targeting different backends from *NumPy* and *PyTorch* in *Python* to *Julia* and *MATLAB* [77]. A similar graph-based approach is taken by *Neuroblox* in *Julia*, combining NMMs with spiking neuron models to build true multiscale models [78].

#### 9.4.2 Performance Optimized Tools

Fast simulations, meaning computationally efficient ones, form the basis for many experiments performed using BNMs, particularly parameter sweeps or optimizations. This need has led to tools that implement critical components, often originating from *Python* tools like TVB, in lower-level languages like *C* [21] or *C++* [22] to optimize performance on modern, highly parallel, hardware including GPUs [79]. Another direction, exemplified by *vbjax* [80], is to target modern libraries like *JAX* [26] to access contemporary hardware and utilize state-of-the-art numerical methods such as automatic differentiation (cf. Section 4.3.2). However, these low-level approaches often achieve maximal performance at the cost of ease of use and comprehensive documentation.

#### 9.4.3 Specialized Tools

The last category loosely groups all the one-off implementation found in the literature. They often implement a single NMM in a novel context or framework, representing the cutting edge of research. Some examples include using a implementation based machine learning framework *PyTorch* to fit fMRI data [25] or evoked potential during transcranial magnetic stimulation [81]. Others like the *Bayesian Virtual Epileptic Patient* [23] leverages bayesian inference for a personalized model of epileptic spread. Expanding on this methodological focus, the *Virtual Brain Inference (VBI)* toolkit [24] offers a more general, flexible framework specifically designed for efficient Bayesian parameter estimation across various virtual brain models and neuroimaging modalities.

While highly valuable as proofs of concept for extending the understanding of brain dynamics, these implementations are often limited in general utility and reusability, and frequently lack comprehensive documentation.

### 9.5 Limitations Bifurcations and Unstable Gradients

Although our approach successfully scales to thousands of parameters, fundamental challenges arise when optimizing across dynamical regimes. Ideally, we would apply multimodal optimization to more biologically detailed models like Jansen-Rit, which captures richer oscillatory dynamics than the simplified Hopf model. However, combining long FC simulations with models exhibiting positive Lyapunov exponents, meaning the system amplifies small perturbations, poses significant challenges for gradient-based optimization.

To investigate these limitations, we return to the Jansen-Rit model, now applying it to FC fitting, a task requiring long simulation times to capture slow fluctuations in functional connectivity. The relationship between simulation length and dynamical stability becomes critical here, as characterized by the Lyapunov time *τ*_*λ*_ (the inverse of the maximum Lyapunov exponent *λ*_*max*_), which determines how quickly nearby trajectories diverge and thus limits the reliable forecast horizon of the system.

Exploring the effect of global coupling *G* on FC for the Jansen-Rit model (cf. Figure 14) reveals both opportunities and challenges. The correlation with target FC peaks around *G* ≈ 15 (cf. Figure 17 panel **c**), precisely where the system undergoes a fundamental change in dynamics. At this coupling value, the average network activity transitions from fixed-point to oscillatory dynamics (panel **a**), accompanied by the maximum Lyapunov exponent switching from negative to positive, indicating a transition from stable to chaotic dynamics.

**Figure 17.**
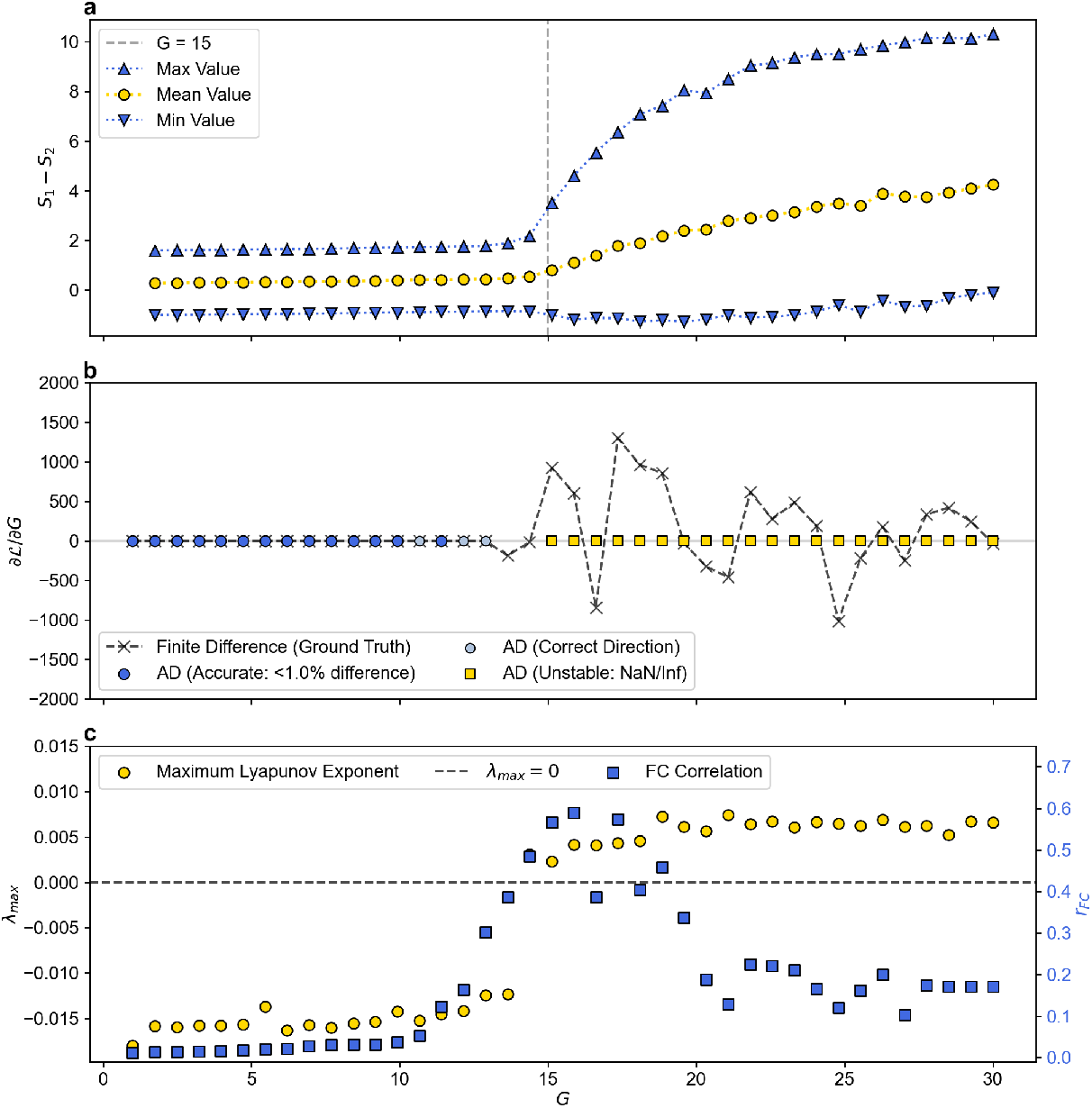
Gradients can turn unstable around bifurcation points. Panel **a** the mean, minimum and maximum of the average network activity for slowly increasing global coupling values *G*. For values around *G* ≈ 15 a bifurcation occurs where the switch from fixed-point to oscillating dynamics occures. In panel **b** the gradient of the FC correlation (*r*_*FC*_) with respect to *G* is displayed. It was obtained with AD and via finite differences as a ground truth, and starts to turn unstable around the bifurcation point. Panel **c** shows a peak in FC correlation around the bifurcation and also shows how the maximum Lyapunov exponent (*λ*_*max*_) switches from negative to being positive.

This bifurcation has profound consequences for gradient-based optimization. Panel **b** shows gradients *∂*ℒ/*∂G* calculated via both AD and finite differences as ground truth. While these agree in the non-oscillating, low-coupling regime where *λ*_*max*_ < 0, the AD-based gradient becomes unstable near the bifurcation point. This instability manifests as gradient values increasing by several orders of magnitude for small parameter changes, quickly leading to NaN values that cause optimizer failure.

The fundamental issue is that when simulation time exceeds *τ*_*λ*_, gradient information propagated through the computational graph accumulates exponential sensitivities. For FC calculations requiring minutes of simulated time, even weakly chaotic dynamics with positive *λ*_*max*_ render gradients unreliable. While specialized techniques exist to handle such cases, including gradient clipping or switching to gradient-free optimization near bifurcations, our current implementation requires monitoring *λ*_*max*_ during optimization to detect and avoid these unstable regimes. This represents a key area for future development in making differentiable brain dynamics robust across dynamical regimes.

#### 9.5.1 Bistability in the Wong-Wang Model

In Section 2.1, we found that optimal fits for FC and FCD using the reduced Wong-Wang model occur near a cusp bifurcation, which creates a bistable regime. While the uncoupled model shows no bifurcation at the best-fit parameters (Figure 12 panels **a** and **b**), the coupled network exhibits bistability (panels **c** and **d**). We identified this bistable behavior by slowly increasing and then decreasing the global coupling parameter *G*, observing hysteresis in the average network activity as it switches between low and high activity states.

Near such bifurcations, small parameter changes induce large dynamical shifts, resulting in extremely large gradients. Combined with floating-point precision limitations, this can lead to exploding or vanishing gradients, a well-known problem in deep recurrent neural networks [47]. Even without catastrophic failure, these large gradient variations can destabilize optimization or slow convergence.

#### 9.5.2 Gradient Clipping

One simple solution to address gradient instabilities near bifurcations is through gradient clipping [47], which bounds gradient magnitudes to stabilize optimization near these steep regions of the loss landscape. Applying this technique in Section 2.1 resulted in more reliable convergence and fewer local optima. The clipping threshold must be chosen carefully: too low and optimization progresses slowly, too high and instabilities persist.

#### 9.5.3 Gradient Flossing

An alternative approach is to include the Lyapunov exponent itself as a differentiable constraint within the loss function. This approach, termed *gradient flossing*, has successfully stabilized training in recurrent neural networks while improving performance [48]. The method works by penalizing positive Lyapunov exponents, effectively constraining the optimization to dynamical regimes where gradients remain stable.

However, unlike neural network weights which lack direct physical interpretation, BNM parameters carry more direct physiological meaning. Each parameter often corresponds to specific biophysical properties (time constants, coupling strengths, etc.), making it unclear which parameters should be constrained through gradient flossing. Constraining the wrong parameters could force the model away from physiologically realistic regimes while satisfying the Lyapunov constraint.

#### 9.5.4 Shadowing Methods

Conversely, some physiological models may require chaotic dynamics [82], where positive Lyapunov exponents are not a bug but a feature. For these cases, shadowing methods based on the shadowing theorem offer a path forward [49]. Despite exponential divergence in chaotic systems, the theorem guarantees that a true trajectory exists near any numerically computed trajectory. By finding these shadow trajectories, we can compute meaningful gradients of time-averaged quantities even in chaotic regimes.

However, two key limitations constrain their applicability. First, these methods require special adaptations for systems with delayed states [50], which are ubiquitous in brain network models due to axonal transmission delays. Second, their computational cost scales poorly with network size, limiting practical application to smaller models. The computational complexity grows rapidly as the number of degrees of freedom increases, making shadowing methods currently impractical for large-scale brain networks with hundreds of regions.

#### 9.5.5 Implementation: Delay Parameter Optimization

The current implementation uses fixed step size integration, which requires collecting delayed states by directly indexing into a history array. This indexing operation has non-smooth derivatives, preventing gradient-based optimization of parameters such as conduction speed or time delays. A solution would be to replace the discrete history array with a continuous, differentiable interpolation function that can be evaluated at arbitrary time delays *τ* . This modification would enable gradient-based optimization of delay parameters while also supporting adaptive time step integration, improving both stability and computational efficiency for non-stiff systems [51].

### 9.6 From Point Estimates to Parameter Distributions

While gradient descent powerfully optimizes brain network models, the gradient information itself enables a much broader range of inference approaches. Once gradients are available, researchers can seamlessly transition from point estimates to full distributional analyses using the same underlying computational infrastructure, as scientific understanding often requires more than a single best parameter set.

Researchers frequently need insights into parameter distributions and their uncertainties, naturally leading to Bayesian approaches where parameter distributions are integral to the modeling framework. However, posterior distributions in complex models quickly become intractable, necessitating approximation methods that can leverage the same gradient computations used for optimization.

Variational inference exemplifies this synergy by approximating intractable posteriors with simpler distributions, minimizing their difference through the Kullback-Leibler divergence. Since this divergence cannot be minimized directly, practitioners instead maximize the Evidence Lower Bound, effectively transforming Bayesian inference into the same gradient-based optimization framework already established for point estimation [52]. The gradient infrastructure exemplified throughout this paper thus directly enables sophisticated uncertainty quantification.

For more accurate posterior sampling, Markov Chain Monte Carlo methods like Hamiltonian Monte Carlo leverage the same gradient information to efficiently explore complex distributions [53,83]. While these methods provide asymptotically exact samples, they require hundreds of gradient evaluations per sample, quickly exceeding the computational cost of finding point estimates through direct optimization. Their effective use demands substantial domain expertise and should be reserved for cases where uncertainty quantification justifies the computational expense. They work particularly well as refinement steps after initial optimization provides good starting points.

When gradient instabilities prevent these gradient-dependent approaches (cf. Section 9.5), gradient-free alternatives become essential. Simulation-based inference addresses this by generating large datasets from prior parameter distributions, then training neural density estimators to approximate posteriors given empirical observations [42,84]. This approach bypasses gradient computation entirely while maintaining the benefits of Bayesian inference.

TVB-Optim’s flexibility and capability to leverage distributed computing facilitates all these approaches, enabling researchers to transition seamlessly with a single model definition from point estimates through variational approximations to full Monte Carlo sampling as their scientific questions demand. The gradient infrastructure established for optimization becomes the foundation for this entire spectrum of inference methods.

## Glossary

*AD*: Automatic Differentiation
*BNM*: Brain Network Model
*BOLD*: Blood-Oxygenation-Level–Dependent imaging
*EEG*: Electroencephalography
*FC*: Functional Connectivity
*FCD*: Functional Connectivity Dynamics
*FIC*: Feedback Inhibition Control
*KS*: Kolmogorov-Smirnov
*MEG*: Magnetoencephalography
*ML*: Machine Learning
*NMM*: Neural Mass Model
*SDDE*: Stochastic Delay Differential Equation
*SGD*: Stochastic Gradient Descent
*TVB*: The Virtual Brain
*TVB-O*: The Virtual Brain Ontology
*fMRI*: Functional Magnetic Resonance Imaging

